# Rapid protein evolution by few-shot learning with a protein language model

**DOI:** 10.1101/2024.07.17.604015

**Authors:** Kaiyi Jiang, Zhaoqing Yan, Matteo Di Bernardo, Samantha R. Sgrizzi, Lukas Villiger, Alisan Kayabolen, Byungji Kim, Josephine K. Carscadden, Masahiro Hiraizumi, Hiroshi Nishimasu, Jonathan S. Gootenberg, Omar O. Abudayyeh

## Abstract

Directed evolution of proteins is critical for applications in basic biological research, therapeutics, diagnostics, and sustainability. However, directed evolution methods are labor intensive, cannot efficiently optimize over multiple protein properties, and are often trapped by local maxima. *In silico-*directed evolution methods incorporating protein language models (PLMs) have the potential to accelerate this engineering process, but current approaches fail to generalize across diverse protein families. We introduce EVOLVEpro, a few-shot active learning framework to rapidly improve protein activity using a combination of PLMs and protein activity predictors, achieving improved activity with as few as four rounds of evolution. EVOLVEpro substantially enhances the efficiency and effectiveness of *in silico* protein evolution, surpassing current state-of-the-art methods and yielding proteins with up to 100-fold improvement of desired properties. We showcase EVOLVEpro for five proteins across three applications: T7 RNA polymerase for RNA production, a miniature CRISPR nuclease, a prime editor, and an integrase for genome editing, and a monoclonal antibody for epitope binding. These results demonstrate the advantages of few-shot active learning with small amounts of experimental data over zero-shot predictions. EVOLVEpro paves the way for broader applications of AI-guided protein engineering in biology and medicine.

## Introduction

Protein diversity has been shaped by billions of years of evolutionary pressures, filtering potential design space for diverse biological activities. Emerging evidence suggests that these sequences embody a fundamental language of biology that can be modeled with deep learning to offer unique insights into the evolutionary processes that have sculpted life on our planet. Protein language models (PLMs) learn the grammar of protein diversity by training to complete masked amino acids across comprehensive protein sequence databases, generating emergent biological representations(*1–4*). Leveraging this rich representation of the evolutionary landscape, PLMs have been used to nominate protein variants with improved activity(*5*, *6*). However, these approaches have had limited success. Generative PLMs, such as ESM3, ProtGPT2, and ProGen(*7–9*), can design novel proteins, but these de novo-designed variants typically only reach wild-type level activity after extensive rounds of experimental testing(*7*, *10*). This inability of PLMs to substantially improve protein activity in zero-shot settings is driven by their inability to generalize to new contexts due to evolutionary constraints and limited training data. Active learning methods that combine context-specific data with deep learning models, including machine learning-directed protein evolution (MLDE) methods(*11–13*), have effectively improved diverse proteins(*14*, *15*) but at the cost of comprehensive experimental evaluation. Merging active learning with PLMs may simplify the evolution process and overcome this shortcoming, but previous attempts (*16*) have not generalized well beyond proof-of-concept demonstrations like fluorescent protein engineering.

Here we present a novel protein evolution model, EVOLVEpro (**EVO**lution **V**ia **L**anguage model-guided **V**ariance **E**xploration for **pro**teins). EVOLVEpro evolves high-activity protein variants with few-shot learning and minimal experimental testing, achieving accurate prediction of high-activity mutants from sequence alone. This performance stems from a modular approach, combining an evolutionary-scale PLM with a top-layer regression model to learn a protein’s activity landscape and guide the directed evolution process *in silico*. By applying EVOLVEpro in a few-shot active learning framework, protein sequences with significantly higher activity can be nominated in a generalizable fashion with minimal effort. The modularity of the EVOLVEpro architecture allows this framework to scale with larger parameter PLMs. Moreover, EVOLVEpro prompting only uses protein sequences and does not require structural information, expert knowledge, or prior data. We demonstrate EVOLVEpro’s ability to evolve multiple activities of a protein simultaneously, opening up vast possibilities for its use in biology and medicine.

We benchmark EVOLVEpro *in silico* across a panel of 12 different protein datasets, showing state-of-the-art performance, and then apply the final model for three different applications (antibody drugs, genome editing, and mRNA manufacturing) with five proteins: 1) a monoclonal COVID antibody, 2) a miniature CRISPR nuclease, 3) a Bxb1 integrase, 4) a prime editor, and 5) a T7 RNA polymerase. This demonstration is, to the best of our knowledge, the first demonstration of PLM-guided models to demonstrate utility in evolution across diverse protein families in non-reporter proteins like GFP. EVOLVEpro yields mutants with 2- to 515-fold improvement over initial proteins. We demonstrate *in vivo* liver editing with an EVOLVEpro-engineered miniature nuclease and improved *in vivo* mRNA performance generated from an EVOLVEpro evolved T7 polymerase. Analyzing nominated mutations with the structure of the protein, EVOLVEpro explores diverse residues that are often in non-intuitive locations of the protein. Moreover, we found that the learned activity landscape is distinct, and often negatively correlated with, the fitness landscape inferred by the underlying protein language model. Lastly, we showcase EVOLVEpro’s utility in proposing multi-mutant protein sequences out of a vast sequence space that enables final mutants with much higher activity than naturally observed proteins. EVOLVEpro establishes the capabilities of few-shot active learning with protein language models for optimizing proteins for diverse activities.

## Results

### Development and benchmarking of the EVOLVEpro model

We developed a deep learning based directed evolution framework called EVOLVEpro that involves: 1) a PLM to encode protein sequences into a continuous latent space to facilitate activity optimization, and 2) a top-layer regression model to learn the mapping between latent space and activity from a few number of data points (i.e. the low-N regime). We employ active learning by using the regression model to rank protein sequences according to their predicted fitness from which we select the top-ranked sequences to experimentally validate. This cycle is performed iteratively to evolve defined protein activities until they reach desired levels (Fig. 1A).

**Fig. 1.**
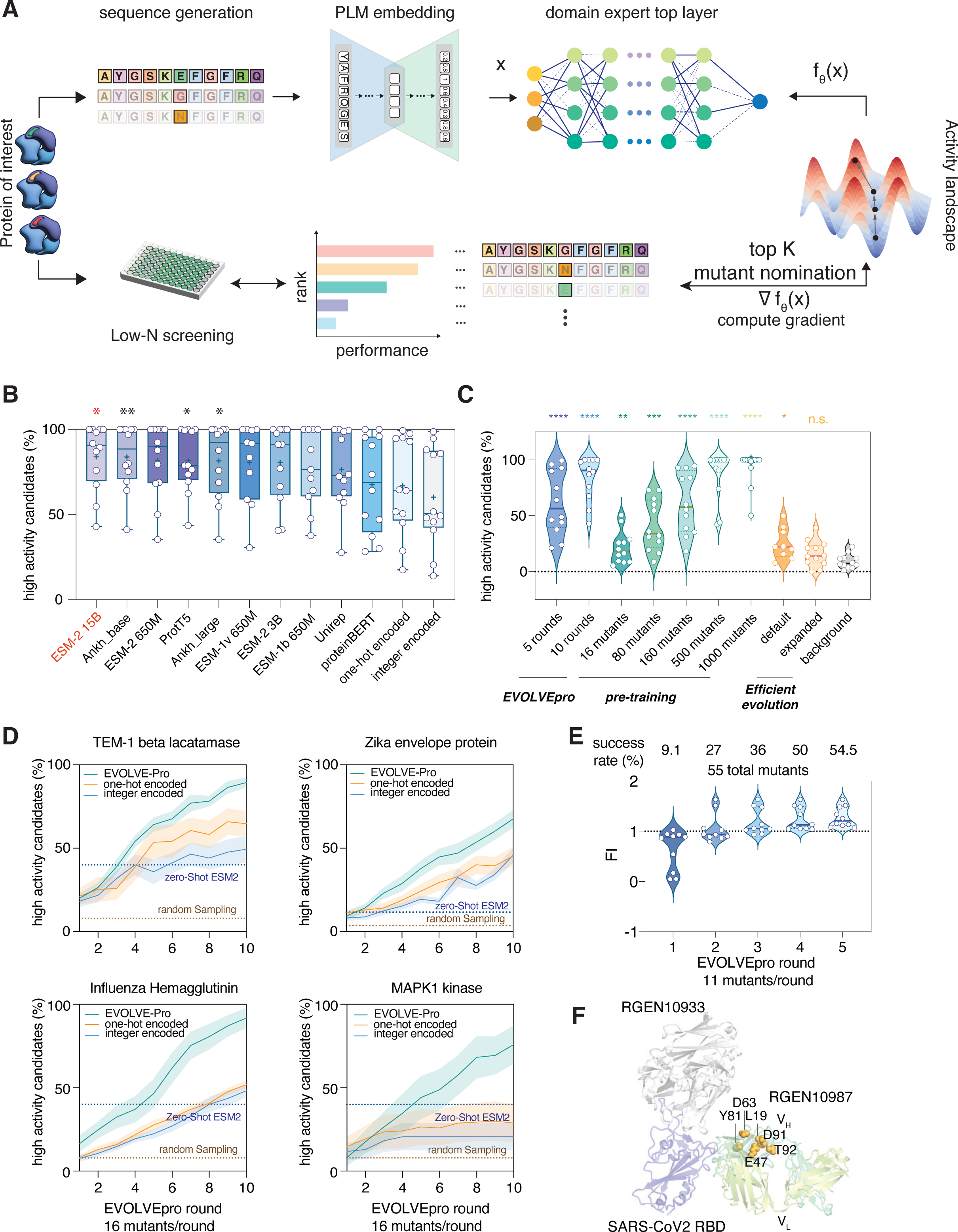
Developing and benchmarking EVOLVEpro for protein language model-guided engineering. (**A**) Schematic describing the EVOLVEpro method. Proteins of interest go through iterative rounds of low-N screening. A foundational PLM generates embeddings for all mutants of a protein and the average embedding by pooling across all residues is used as input for the top-layer model. Each mutant’s activity is experimentally determined and used to train a domain expert top-layer model with PLM embedding as input. The top-layer model then nominates the top-N mutants for the next round of testing and the weights are updated iteratively in an active learning format. (**B**) Benchmarking of foundational models across a panel of 12 comprehensive deep mutational scanning (DMS) datasets. Each point is a unique protein and its DMS data. ESM2-15B has the highest average percent success in high activity variants prediction. (**C**) Comparison between EVOLVEpro in active learning format, in zero-shot pretraining format, and an existing zero-shot prediction method using protein language model (*6*) across 12 DMS datasets. Each point is a unique protein using its DMS data. (**D**) Performance over 10 rounds of EVOLVEpro with 16 mutants per round, compared to two different non-language model encoding schemes (one-hot encoding and integer encoding). Model performance is benchmarked on four datasets(*17*, *22*, *26*, *54*) and compared to zero-shot ESM2 nomination success rate and background random sampling (*6*). Error bar represents the standard deviation for n=10 random simulations. (**E**) Engineering of REGN10987 over five rounds of EVOLVEpro. Data shows cumulative top 10 mutants’ fold improvement over wild-type binding affinity to the target antigen across 5 evolution rounds. Percentages show the percent of mutants that have higher activity than wild-type REGN10987 each round. (**F**) Mapping of the top mutations on the structure of REGN10987 (PDB: 6XDG).

We optimized EVOLVEpro across five parameters: 1) the strategy employed for the first round mutant selection, 2) the top-layer regression model that learns the activity landscape, 3) the active learning strategy for selecting mutants for the next round, 4) data processing for experimentally measured activities, and 5) the PLM embedding vector transformation (table S1, Data S1). To perform a grid search across this space, we curated a panel of twelve unique deep mutagenesis scanning (DMS) datasets for *in silico* validation (*17–27*) (table S2, Data S2). These twelve proteins represent diverse activities, including viral spike proteins, RNA-guided nucleases, lactases, and kinases, ensuring that the resulting model will be generalizable for learning diverse protein activity landscapes in the PLM latent space.

We first focused on the ESM-2 protein language model because of its large training data and available model size of >200M proteins and 15B parameters, respectively. Using the ESM-2 15B parameter model, our grid search found the optimal strategy was: 1) selecting a random set of first-round variants, 2) employing a random forest regressor discriminatory model to predict protein activities, 3) using residue pooled average embeddings, and 4) using a top-N selection strategy in each round of evolution (fig. S1A). This model nominated a high frequency of gain-of-function protein variants in only 5 rounds (fig. S1A-B). Since we focused on the percent of variants passing an activity threshold as our evaluation metric in the grid search, we next checked for increasing activity during *in silico* evolution. We found that both the median activity and the activity of the nominated top mutant increased monotonically from round to round across all DMS datasets, further validating the model’s performance in this low-N active learning setting (fig. S1B).

In general, 16 mutants per round of evolution for 10 rounds identified top mutants with fitness up to 7-fold higher activity than starting wild-type sequence (fig. S1B). To understand how the number of variants per round affected performance, we tested between 10 and 100 variants per round, finding that larger rounds increased prediction accuracy (fig. S1C). This performance trade-off indicates that EVOLVEpro can be used for both extremely low-N evolution (<20 mutants per round) for rapid and cheap experimental characterization and medium-N (∼100 mutants per round) for quicker and more efficient evolution with fewer rounds.

After optimizing the top-layer model and learning strategies, we optimized the PLM, comparing ESM-2 15B to a panel of foundational models. Using the optimal parameters from the grid search, we benchmarked performance against smaller versions of ESM-2 and ESM-1(*28*), UniRep(*16*, *29*), ProtT5(*30*), ProteinBERT(*4*), Ankh(*3*), one-hot encoding, and integer encoded protein representations. ESM-2 15B parameter model outperformed all the other models for identifying the highest fitness proteins for all datasets except two, confirming its final selection for the EVOLVEpro latent space model (Fig. 1B, Data S3). Importantly, large parameter PLMs showed a significant boost in prediction accuracy compared to non-language model-based architectures, indicative of the powerful feature extraction present in transformer-based models (Fig. 1B).

We next benchmarked EVOLVEpro’s performance relative to other PLM-based engineering approaches. As many methods require pre-training a discriminatory model on thousands of variants, we tested versions of EVOLVEpro augmented with various amounts of pre-training (Fig. 1C). Active learning drastically reduced the overall number of mutants required: EVOLVEpro with only 5 rounds of evolution (16 mutants per round) was equivalent in performance to EVOLVEpro pre-trained with 160 mutants, while 10 rounds of evolution (16 mutants per round) was equivalent to pre-training with 500 mutants. Moreover, EVOLVEpro significantly outperformed zero-shot prediction methods (*6*). This comparison confirms that the few-shot nature of EVOLVEpro allows for efficient directed evolution with minimal effort and low-N testing per round (Fig. 1C, Data S4).

Lastly, we analyzed the per-round evolution improvement for EVOLVEpro compared to one-hot and integer encoding and zero-shot prediction, finding that by round 5, variants with significantly enhanced fitness could universally be found (at 16 mutations per round) (Fig. 1D, fig. S2). Moreover, in many cases, the one-hot and integer encoding frameworks saturated much earlier in the evolution process and never reached the fitness levels achieved by EVOLVEpro. Interestingly, for some proteins we observe a non-linear increase in protein fitness after round 3, suggesting greater gains in mapping the protein fitness landscape as EVOLVEpro evolution proceeds.

### Antibody optimization with EVOLVEpro

As a first test of EVOLVEpro, we optimized the binding interaction of the REGN10987 antibody to the extracellular epitope of the SARS-CoV-2 spike protein. REGN10987, a component of an approved COVID-19 therapy (*31*), is engineered for neutralization of the SARS-CoV-2 spike protein and is refractory to previous *in silico* optimization (*6*). Given the challenge of improving REGN10987, it is an ideal first test of EVOLVEpro’s capabilities. Mutagenizing the heavy chain variable region with EVOLVEpro, we compared wild-type REGN10987 to 11 nominated mutants each round with an enzyme-linked immunosorbent assay (ELISA) against the SP6 stabilizing variants of SARS-CoV-2 spike protein(*32*). We found significant fold improvement (FI) in binding from only 1 round of EVOLVEpro, and five rounds of mutagenesis yielded variants that showed a 63% improvement by ELISA with an IC_50_ of 11.9 nM (Fig. 1E, fig. S3A). We observed that the success rate of EVOLVEpro increased each round, with success increasing from 9.1% in the first random round to 54.5% in the last round. This improvement demonstrates EVOLVEpro’s top-layer model learns an activity grammar distinct from the fitness grammar captured initially by the underlying foundational model (Fig. 1E, fig. S3B).

In the structure of REGN10987 bound to the spike epitope, one high-performing mutation (D63K) lies inside the 2nd complementarity-determining region (CDR), with other mutations clustered around the CDR3 in the framework region (HFR3), suggesting a role in enhanced antigen binding (Fig. 1F). To understand EVOLVEpro’s mutational trajectory, we represented the model’s attention to particular residues as cumulative frequency of individual residues being explored by the model and found that multiple residues are repeatedly explored by the model including D63, D91, and N32 (fig. S3C). Across all nominated mutations, successive rounds of training focused on regions around CDR2, emphasizing the increased attention of the EVOLVEpro to specific regions of the protein. We next analyzed each mutant’s observed activity versus the PLM-predicted fitness. We calculated the mutant fitness as predicted marginal masked score within the ESM2 embeddings (pMMS) and found that the activities of EVOLVEpro variants did not correlate with fitness (fig. S3D). To extrapolate this finding across the entire REGN10987 mutational landscape, we projected the base layer PLM fitness score and top-layer random forest predicted fold improvement (pFI) in the latent space for every possible single mutation variant (fig. S3E-F). There was relatively little overlap between the two distributions, with a negative correlation of −0.22 between predicted fitness and predicted activity (fig. S3G).

### Evolution of a miniature RNA-guided CRISPR nuclease with EVOLVEpro

Programmable RNA-guided nucleases have diverse applications in basic biology, therapeutics, and diagnostics. However, commonly used nucleases, such as the Cas9 from *Streptococcus pyogenes* (SpCas9) are too large to effectively be packaged in common viral vectors such as adeno-associated viral (AAV), and more compact high-efficiency nucleases, such as the Cas9 from *Staphylococcus aureus* (SaCas9) still preclude the use of larger regulatory elements or protein fusions. Miniature Cas12f nucleases have compact sizes (<700 residues) but suffer from reduced efficiencies, requiring significant engineering for genome editing applications(*33*). Previous Cas12f engineering efforts relied on DMS or rationally designed mutations to increase the *in vitro* cleavage activity(*19*, *34–37*), requiring extensive screening to find the optimal variant. To accelerate miniature nuclease engineering, we tested whether EVOLVEpro could rapidly develop highly active Cas12f variants.

We selected the Cas12f from *Pseudomonas aeruginosa* (PsaCas12f) for evolution with set indel formation at the endogenous *RNF2* locus target site as the optimization metric (Fig. 2A). After four rounds of evolution of 12 single mutants per round, EVOLVEpro yielded point-mutants of PsaCas12f with up to 4.9-fold improvement in indel formation. This top variant, PsaCas12f ^K333V^, had >40% indel efficiency at the *RNF2* site (Fig. 2B, fig. S4A). To identify synergies between EVOLVEpro nominated mutants, we combined the top-performing variants from previous rounds in a fifth round. We evaluated a set of these multi-mutants and found that PsaCas12f ^I178A/K333V/K454P^ exhibited greater than 50% indel activity at the *RNF2* locus, a 25% higher activity than any of the single mutants (fig. S4A). Given its performance, we refer to the PsaCas12f ^I178A/K333V/K454P^ variant as EVOLVEpro PsaCas12f (epPsaCas12f). Because these mutants synergize to produce an even more active enzyme, it suggests that the variants identified by EVOLVEpro are uniquely independent in mechanism, highlighting the insightful potential of the method.

**Fig. 2.**
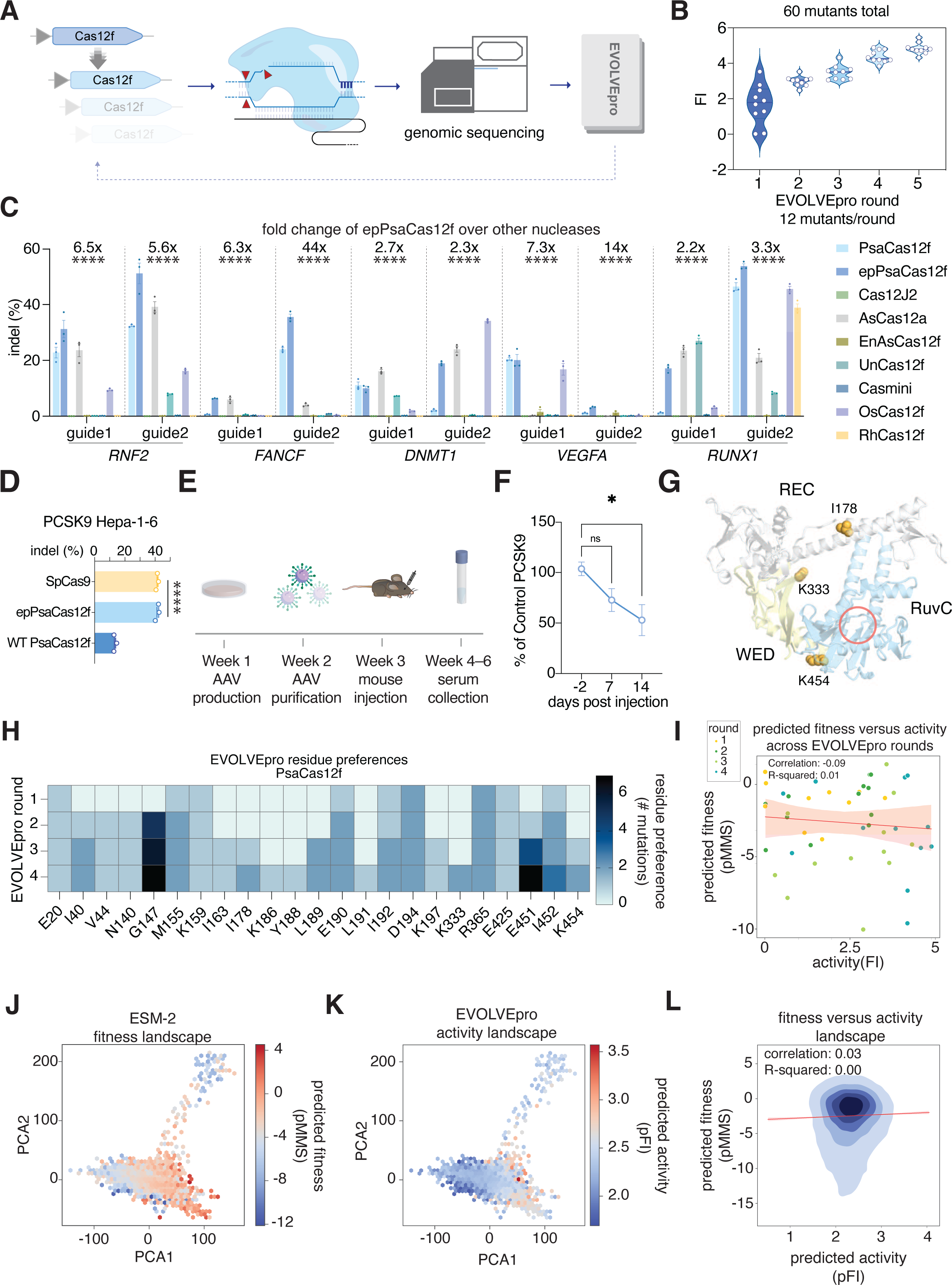
Evolution of highly active miniature CRISPR nucleases with EVOLVEpro. (**A**) Schematic of the evolution strategy with EVOLVEpro for engineering a miniature Cas12f. (**B**) Engineering of PsaCas12f over four rounds of EVOLVEpro and a rational combination multi-mutant round. Data shows cumulative top 10 mutants from current and preceding rounds, as measured by fold improvement of indel activity at the endogenous RNF2 genomic locus. (**C**) Indel activities of WT PsaCas12f, epPsaCas12f, and a panel of published Cas12a and Cas12f nucleases on 10 different genomic targets across five genes (two guides per gene). The fold change on top of each guide denotes the relative fold increase of epPsaCas12f compared to the average of the other published Cas12a and Cas12f nucleases. A one-way ANOVA is performed for each guide sequence shown (****, p<0.0001). (**D**) Next-generation sequencing quantified indel formation at murine PCSK9 genomic loci by epPsaCas12f, WT PsaCas12f, and SpCas9. A one-way ANOVA is performed for each guide sequence shown (****, p<0.0001). (**E**) Schematic of the *in vivo* validation assay for EnPsaCas12f editing at the murine PCSK9 locus for PCSK9 reduction. (**F**) Serum PCSK9 levels at three different time points from −2 days of injection to +14 days. The percent of control PCSK9 was calculated by normalizing to the control group with PBS injected. A two-sided Student’s t-test was run on each time point relative to −2 days’ baseline PCSK9 level (ns, non-significant, *, p<0.05). (**G**) Mapping of the top mutations on the AlphaFold3 model of PsaCas12f. The RuvC active site is indicated by a red circle. (**H**) Heatmap showing most common PsaCas12f mutations explored by EVOLVEpro over rounds of evolution. Any position explored more than once is shown on a cumulative basis across rounds. (**I**) Scatter plot comparing the predicted naive ESM-2 protein fitness (predicted masked marginal score) and scaled tested activity of nominated mutants across evolution, scatter points are colored by rounds in evolution. (**J-K**) Comparison of the PsaCas12f embedding latent space with either predicted naive ESM-2 protein fitness landscape or EVOLVEpro protein activity landscape. (**L**) A kernel density estimate plot of protein fitness as predicted by ESM-2 versus protein activity as predicted by EVOLVEpro. The correlation and linear regression line are shown in red and the R square of the correlation is reported.

To generalize epPsaCas12f’s improved activity, we evaluated the enzyme at 10 different targets across five endogenous genomic loci, comparing to WT PsaCas12f and seven previously characterized Cas12 effectors, AsCas12a, Cas12Φ, UnCas12f1, enAsCas12f, OsCas12f, RhCas12f, and CasMINI(*19*, *35*, *37–40*). We observed consistently higher epPsaCas12f activity compared to WT PsaCas12f on 9 of 10 tested targets (Fig. 2C). Moreover, epPsaCas12f edited the 10 targets with a 23.3 ± 16.7% average indel rate, surpassing all tested miniature Cas12f effectors and AsCas12a with 2.2- to 44-fold improvement. Interestingly, epPsaCas12f generated an average deletion of 5-bp across the 10 tested targets, larger than the deletions generated by other orthologs (fig. S4B). Consistent with our mammalian data, purified epPsaCas12f exhibited higher biochemical DNA cleavage activity than WT PsaCas12f (fig. S4C). Together, these data demonstrate that epPsaCas12f is a highly active, compact effector for mammalian genome editing that outperforms other small effectors.

We applied epPsaCas12f for *in vivo* genome editing applications, using its compact size for single-vector viral delivery *in vivo*. We designed guides targeting a sequence 5′ of exon 3 in the mouse *PCSK9* gene (Fig. 2D). The PCSK9 protein regulates blood low-density lipoprotein (LDL) by binding to LDL receptors, making it a valuable therapeutic target(*41*). We first tested the efficacy of epPsaCas12f in a murine hepatocyte cell line (Hepa 1-6) by co-transfecting murine codon-optimized epPsaCas12f and sgRNA targeting sequences 5′ of exon3 in the *PCSK9* gene. Analyses of epPsaCas12f, WT PsaCas12f, and *Staphylococcus pyogenes* Cas9 (SpCas9) revealed that epPsaCas12f robustly edited *PCSK9* in Hepa1-6 cells with ∼40% indel formation (comparable levels to SpCas9 and 3-fold higher than the WT PsaCas12f) (Fig. 2D).

After validation of epPsaCas12f in Hepa1-6 cells, we packaged both epPsaCas12f and its sgRNA targeting *PCSK9* in a single AAV2/8 vector (Fig. 2E). AAV-epPsaCas12f was administered at a titer of 1.5✕10^12^ viral genome copies per mouse via retro-orbital injection into 3-month-old C57BL/6J mice. We tracked blood PCSK9 levels for 14 days post-injection of AAV and found a significant decrease to around 50% of the original levels after 14 days (Fig. 2F, fig. S4D). We then harvested the liver at day 15, isolated the genomic DNA, and performed next-generation sequencing to survey for indel formation at the *PCSK9* target site (fig. S4E-F). We found around 7% on-target indel formation in the AAV-epPsaCas12f injected mice (fig. S4E), demonstrating that epPsaCas12f can be used for single-vector AAV-mediated genome editing. To survey off-targets, we used Cas-OFFinder to predict the top four off-target cleavage sites generated by epPsaCas12f and analyzed the guide-dependent off-target cleavage in the liver(*42*). We only found detectable editing at one of the four sites with a maximum level of 0.27% indels, confirming minimal off-target cleavage triggered by epPsaCas12f (fig. S4G).

To understand the mechanisms of the beneficial mutations nominated by the EVOLVEpro, we used Alphafold3 to predict the structure of PsaCas12f (Fig. 2G). The predicted structure provides insights into how the PLM-nominated mutations, including I178A/K333V/K454P, contribute to enhancing the DNA cleavage activity (Fig. 2G). The K333V mutation is located in the WED domain, suggesting that it could increase the binding to its RNA guide. The I178A mutation is located in the middle of the long α-helix in the REC domain and forms a hydrophobic core with I245 and L248 in the adjacent α-helix. Given that alanine is a helix-forming residue, the I178A mutation may stabilize the α-helix in the REC domain and thus augment the cleavage activity. The K454P mutation is located at the C-terminus of an α-helix in the RuvC domain and forms hydrophobic interactions with A509 and V511 in the adjacent α-helix, suggesting that it also stabilizes the protein conformation.

We then looked at the model’s attention to particular residues in the protein by calculating the cumulative frequency of individual residues explored by the model and found that multiple residues are repeatedly nominated by the model, including G147 and E451 (Fig. 2H). We calculated the pMMS for each nominated mutant to understand the relationship between the base layer PLM’s fitness prediction and the actual measured protein activity (Fig. 2I). We found that there is a weak negative correlation between fitness and activity in PsaCas12’s local context but EVOLVEpro nominated for higher activity mutants toward the later rounds contrary to high fitness mutants recommended by the PLM base layer (Fig. 2I). We then further projected both base layer PLM’s fitness score and top-layer random forest regressor’s activity score in the EMS2 latent space to better understand EVOLVEpro’s global mutational trajectory (Fig. 2J-L). We found a weak positive correlation of 0.03 between fitness and activity, further denoting the necessity of a top-layer discrimination model to properly distinguish between high fitness and high activity (Fig. 2l).

### Engineering improved prime editors with EVOLVEpro

Many molecular tools, such as next-generation genome editing proteins, function as multiple enzymes acting in concert. Prime editing, which uses an RNA-templated reverse transcriptase to programmably install diverse genome edits, is the fusion of a SpCas9 nicking mutant (nCas9) with an engineered Moloney Murine Leukemia Virus Reverse Transcriptase (M-MLV RT) [D200N, L603W, T306K, W313F, T330P] (termed PE2). We surmised that EVOLVEpro could improve upon these rational mutations, given further optimizations discovered on M-MLV RT by other directed evolution approaches(*43*). As PE-based insertion has difficulty installing longer (>40 nt) edits, we focused on editing outcomes with longer (46 bp) insertions, which have particular utility for programmable gene insertion methods, such as PASTE(*44*). We set up the evolution policy by using a previously described twinPE approach, where two overlapping pegRNAs are used in combination to install a 46bp attB site in the NOLC1 loci in murine hepatocyte cell line (Hepa1-6). Editing was quantified at NOLC1 loci using amplicon sequencing and NGS readout and the top-layer EVOLVEpro model was trained to predict the insertion efficiency (Fig. 3A).

**Fig. 3.**
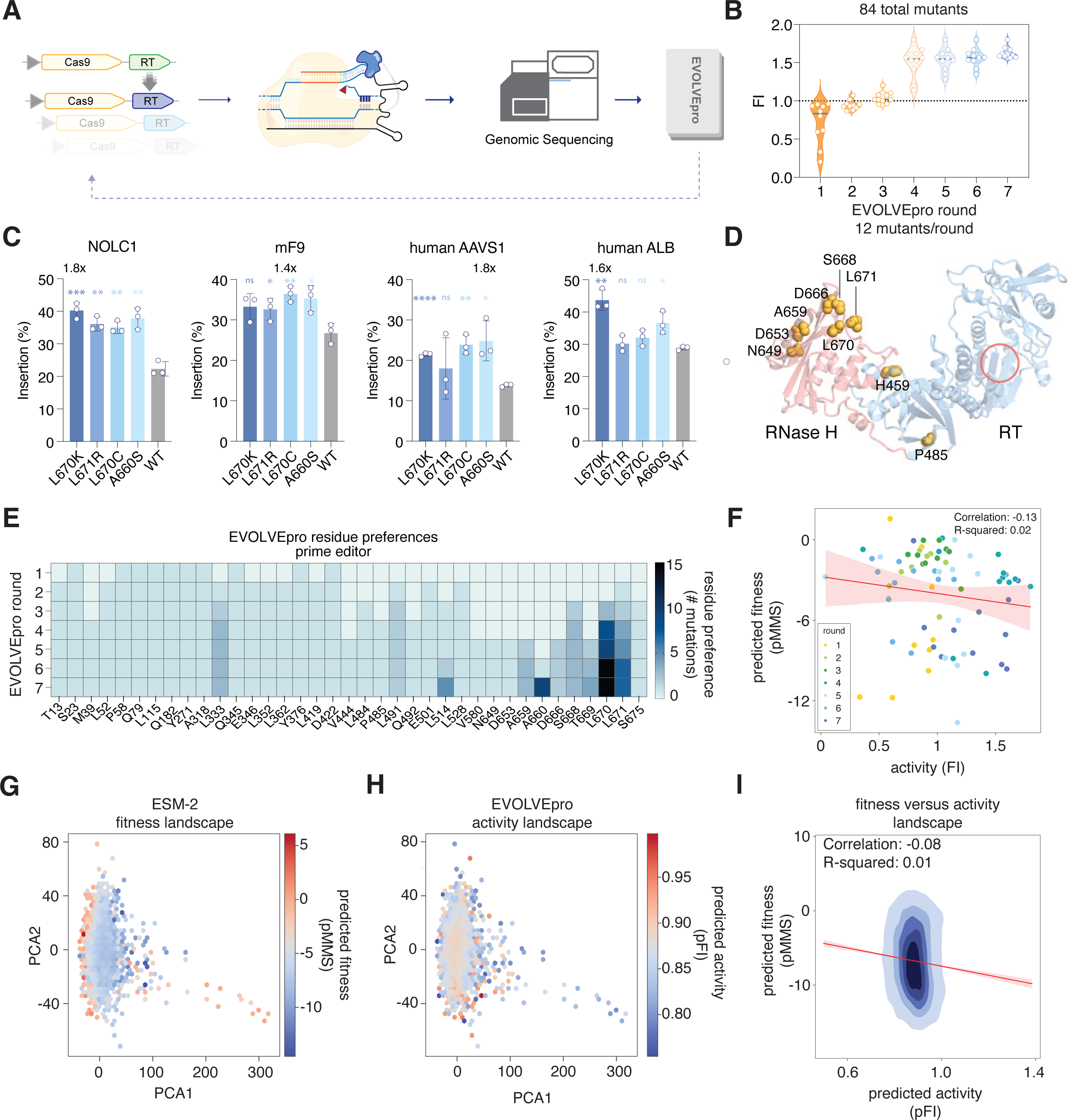
Evolution of prime editor with EVOLVEpro. (**A**) Schematic of the evolution strategy with EVOLVEpro for engineering a prime editor to be more efficient in attB insertion. (**B**) Engineering of the prime editor PE2 with twinPE guides over seven rounds of EVOLVEpro. Data shows cumulative top 10 mutants from current and preceding rounds, as measured by fold improvement of prime editing activity to install a 46 bp AttB site at the murine NOLC1 genomic locus. (**C**) Validation of 4 evolved prime editors in the installation of attB sites at four different endogenous sites in either mouse or human genomes. A two-sided unpaired t-test was run between WT and each evolved prime editor (ns, non-significant, *, p<0.05, **, p<0.01, ***, p<0.001, ****, p<0.0001). Fold change over wild-type PE2 is shown for the best mutant on each genomic locus. Error bars represent standard deviation with n=3 biological replicates. (**D**) Mapping of the top mutations on the AlphaFold3 model of M-MLV RT. The RT active site is indicated by a red circle. (**E**) Heatmap showing most common PE2 mutations explored by EVOLVEpro over rounds of evolution. Any position explored more than once is shown on a cumulative basis across rounds. (**F**) Scatter plot comparing the predicted naive ESM-2 protein fitness (predicted masked marginal score) and scaled tested activity of nominated mutants across evolution, scatter points are colored by rounds in evolution. (**G-H**) Comparison of the PE2 embedding latent space with either predicted naive ESM-2 protein fitness landscape or EVOLVEpro protein activity landscape. (**I**) A kernel density estimate plot of protein fitness as predicted by ESM-2 versus protein activity as predicted by EVOLVEpro. The correlation and linear regression line are shown in red and the R square of correlation is reported.

Over successive rounds of optimization, we found that EVOLVEpro progressively learned the activity landscape of the RT of PE2, yielding improved variants after the initial random selection round and substantially improving upon PE2-based editing by round 4 (Fig. 3B, fig. S5). To check for bias against this single loci in the genome, we took the top 4 performing variants (A660S, L670C, L670K, and L671R) and surveyed their editing efficiency at three additional genomic loci (human AAVS1, human ALB, and mouse Factor IX) in two different cell lines and found statistically significant improvements for A660S in all four sites tested (Fig. 3C). These results point to the general protein activity improvement by our model, delivering an additional set of RT mutations specifically for larger edits.

Projecting the top mutations onto the Alphafold3-predicted structure of the RT reveals that most of them are clustered in the C-terminal RNase H domain (Fig. 3D), which is a surprising result since most PE evolution focuses on RT mutagenesis. We hypothesize that these mutations could alter the cleavage of the template DNA in the RNA-DNA heteroduplex by the RNaseH domain(*45*), facilitating the completion of the prime editing reaction, a route that has not been explored by traditional engineering of prime editors. We then further analyzed EVOLVEpro’s residue site preference during evolution and observed significant attention in residues like L670, L671, and A660, suggesting it was learning that these positions could be quite beneficial for improving activity (Fig. 3E). Analysis of predicted fitness (pMMS) scored by the bottom layer PLM again showed a divergence between fitness and activity for the prime editor (Fig. 3F), allowing EVOLVEpro to successfully use the top discrimination layer to navigate to the higher activity variants in later evolution rounds (Fig. 3F).

Lastly, we try to understand the global mutational trajectory by projecting the activity landscape learned by the random forest regressor and base layer ESM2’s protein fitness landscape onto the first two PCAs of the embedding (Fig. 3G-H). We found almost no convergence between the two distributions with a negative correlation of 0.08 (Fig. 3I). This analysis points again to the divergence between the mutational landscape of a protein’s activity and the commonly used fitness landscape learned during a foundational model’s training on all protein sequences.

### Bxb1 integrase evolution with EVOLVEpro

Large serine recombinases (LSRs) are enzymes that facilitate precise DNA rearrangements, making them crucial tools for genome editing. Their ability to recognize specific DNA sequences and catalyze targeted recombination events allows for efficient and accurate modifications of genetic material, which is essential for advanced gene therapy, synthetic biology, and genetic research. We recently developed a gene insertion technology, PASTE, that leverages LSRs, specifically the Bxb1 integrase, for programmable gene insertion in eukaryotic cells(*44*). A limitation of Bxb1 integrase, however, is its activity saturates in the 20-60% range in cells, limiting the overall integration efficiency that can be achieved. We sought to therefore evolve Bxb1 using EVOLVEpro to improve its activity and demonstrate improved gene integration applications with PASTE in cells.

To evolve Bxb1, we designed a simple integration assay in HEK293FT cells that involved the insertion of an AttP-containing DNA plasmid into an AttB target-containing plasmid (Fig. 4A). Integration can be measured by next-generation sequencing, and the evolution policy is designed to optimize this insertion efficiency. We started evolution with a round of 11 random Bxb1 point mutation variants and then over 8 rounds observed progressively increasing activity resulting in mutants with over 2-fold higher activity than wild-type (Fig. 4B, fig. S6A). As Bxb1 is already fairly active, this fold improvement is expected as we reach near-saturating levels of insertion. To validate the top hits from the evolution campaign, we performed a Bxb1 plasmid titration experiment in a separate cell line (Hela cells) and observed up to 4-fold improvement in recombination efficiency under low Bxb1 expression (Fig. 4C). We further validated the top hits by pre-installing attB sites into the genome of HEK293FT cells using lentivirus and then surveyed for integration efficiency of cargo in the genome. We found up to 4-fold improvement by EVOLVEpro’s mutants compared to wild-type (fig. S6B). We termed T166R the final EVOLVEpro Bxb1 candidate as enhanced Bxb1, or epBxb1.

**Fig. 4.**
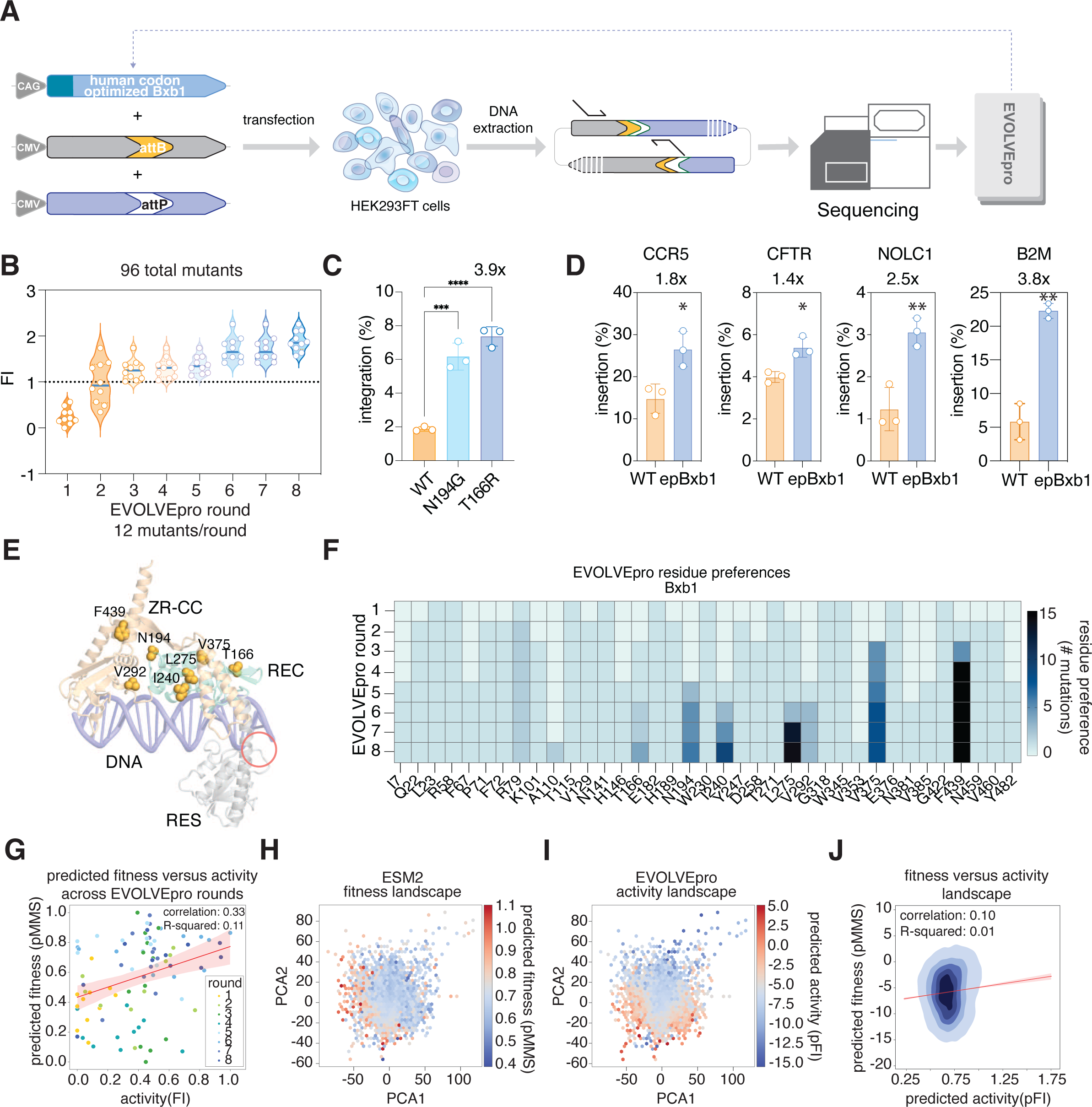
EVOLVEpro engineers enhanced large serine recombinases. (**A**) Schematic of the evolution strategy for evolving the Bxb1 serine integrase from the Mycobacteriophage. (**B**) Engineering of the Bxb1 integrase over 8 rounds of EVOLVEpro. Data shows cumulative top 10 mutants from current and preceding rounds, as measured by fold improvement of plasmid integration over wild-type. (**C**) Performance of top Bxb1 mutants for plasmid recombination with low Bxb1 expression in Hela cell. A two-sided Student’s t-test was run between WT and each evolved Bxb1 integrase (***, p<0.001, ****, p<0.0001). Fold change over wild-type Bxb1 is shown for the best mutant. Error bars represent standard deviation with n=3 biological replicates. (**D**) Validation of epBxb1 with PASTE at four genomic sites across human and mice genomes. A two-sided Student’s t-test was run between WT and each evolved Bxb1 integrase (*, p<0.05, **, p<0.01). Fold change over wild-type Bxb1 integrase is shown for each genomic locus. Error bars represent standard deviation with n=3 biological replicates. (**E**) Mapping of the top mutations on the AlphaFold3 model of the Bxb1 monomer bound to DNA. Bxb1 forms a tetrameric synaptic complex during recombination between two DNA molecules. The active site is indicated by a red circle. (**F**) Heatmap showing most common Bxb1 mutations explored by EVOLVEpro over rounds of evolution. Any position explored more than once is shown on a cumulative basis across rounds. (**G**) Scatter plot comparing the predicted ESM-2 protein fitness score versus experimentally measured bxb1 integration efficiency (scaled) across evolution rounds. The correlation and linear regression line are shown in the plot. (**H-I**) Comparison of the Bxb1 latent space with either predicted ESM-2 protein fitness (masked marginal score) or EVOLVEpro protein activity fold improvement. (**J**) A kernel density estimate of protein fitness as predicted by ESM-2 versus protein activity as predicted by EVOLVEpro. The correlation and linear regression line are shown in red and the R square of correlation is reported.

To test whether the epBxb1 variant’s improved activity can improve the programmable insertion of cargo DNA into the chromosome, we tested this variant in the context of PASTE and compared it against the wild-type Bxb1 across five different genomic loci. We found up to ∼4-fold improvement in the final large cargo insertion rate into the genome, which highlights the generalizable gain in activity (Fig. 4D, fig. S6C).

An AlphaFold3-predicted model of Bxb1 bound to attachment site DNA indicates that the top beneficial EVOLVEpro mutations clustered in the Bxb1 DNA-binding domains, likely increasing the affinity to its DNA targets (Fig. 4E). Of these residues, V292S could directly interact with the phosphate backbone of the target DNA based on its positioning relative to the attachment site, whereas the others likely modulate DNA binding via indirect interactions.

Analysis of the residue exploration by the model revealed that multiple positions, including F439, V375, and L275, are visited up to 15 times; the DNA-interacting residue V292 was also visited multiple times. Overall, this highlights EVOLVEpro’s ability to recognize the functional importance of certain regions in the protein, much like structure-guided engineering approaches (Fig. 4F).

We then calculated the relationship between the fitness (pMMS) and activity (observed fold improvement) for Bxb1 integrase and found a weakly positive correlation between the two metrics contrary to the other proteins reported evolves. This likely reflects a subset of protein families where protein stability and fitness as learned by the PLM can predict activity as previously reported(*46*) (Fig. 4G). However, given that the relationship is weak, a model like EVOLVEpro is still needed to efficiently and quickly reach high-performing variants without encountering many false positives. Lastly, we found that the global mutation landscape learned by EVOLVEpro was still divergent from the predicted fitness (pMMS) by ESM2 with an even weaker correlation, further highlighting the ability of EVOLVEpro to learn protein activity at a global scale and how stability/fitness prediction is not sufficient for rapid and efficient protein evolution (Fig. 4H-J).

### Evolving T7 RNA polymerase for efficient and highly pure RNA production

We finally sought to engineer an enzyme broadly useful for medicine and science, while also demonstrating the versatility of EVOLVEpro to evolve an enzyme in a multi-objective setting. We selected the T7 RNA polymerase (RNAP) due to its critical role in RNA production for mRNA therapies, mRNA vaccines, cell engineering, and basic scientific studies. As mRNA production has numerous features characterizing its potency and quality, as opposed to genome editing where one feature matters the most, we designed a multi-objective optimization function to evolve a high-fidelity T7 RNAP for mRNA production with these three parameters: 1) RNA yield measured via UV-vis spectrophotometry, 2) mRNA translation in a dsRNA sensitive cell line measured via luciferase translation, and 3) RNA purity measured via immunogenicity in BJ fibroblast cells by IFN-beta RNA production (Fig. 5A). We weighted these features in the EVOLVEpro objective function by 20%, 40%, and 40%, respectively to prioritize the higher fidelity and lower immunogenicity aspects of this enzyme for clinical applications. To facilitate high throughput variant testing, we relied on SP6 in vitro transcription-translation coupled reaction kits to generate mutant T7 RNAP in a one-pot reaction and subsequently use the produced T7 RNAP to produce co-transcriptionally capped Cypridina luciferase mRNA for downstream *in vitro* testing.

**Fig. 5.**
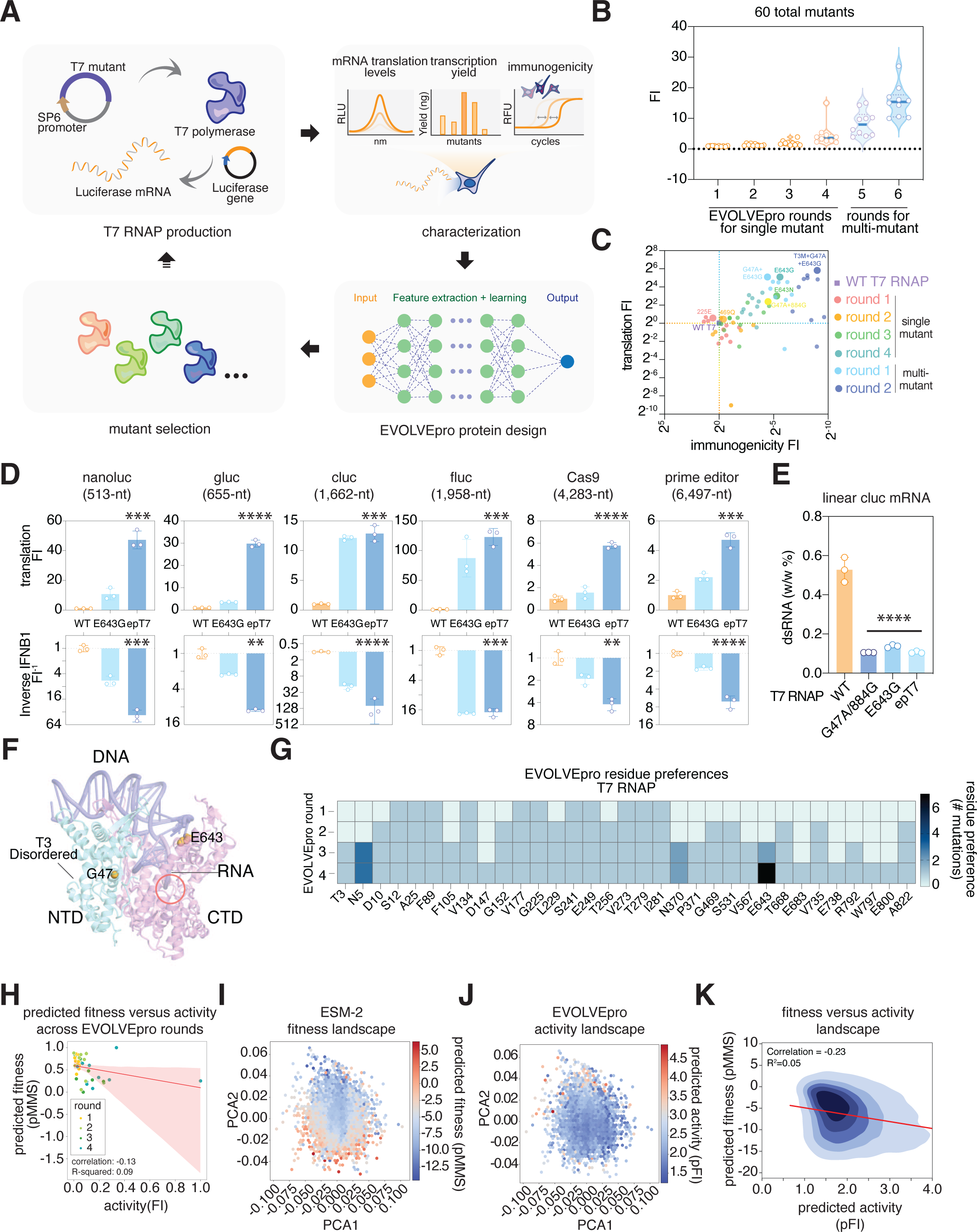
Engineering RNA polymerases for high yield and low immunogenicity mRNA production. (**A**) Schematic of the strategy for high throughput T7 RNA polymerases mutant testing and evolution policy setup for evolving a high fidelity T7 RNAP. (**B**) Engineering of T7 RNAP over six rounds of EVOLVEpro. Data shows the top 10 mutants from current and preceding rounds, as measured by fold improvement of transcription fidelity over wild-type. (**C**) Performance of T7 mutants from six EVOLVEpro rounds and previously engineered G47A/884G SOTA T7 RNAP in Cluc mRNA translation and immunogenicity in BJ Fibroblast cells. (**D**) Validation of epT7 for production of 6 mRNA sequences ranging from 513nt to 6496nt. Purified WT or mutant RNAP is used to produce these sequences, and they were transfected into BJ fibroblast cells for either protein translation readout or targeted IFNB1 gene expression analysis using qPCR 24 hours after transfection. A two-sided Student’s t-test was run between WT and each evolved T7 RNAP (**, p<0.01, ***, p<0.001, ****, p<0.0001). Error bars represent standard deviation with n=3 biological replicates. (**E**) dsRNA ELISA is used to analyze the amount of dsRNA during transcription of a 1662 nt Cypridina luciferase mRNA. 500 ng of post-transcription product is used as input for the dsRNA ELISA. A two-sided Student’s t-test was run between WT and each evolved T7 RNAP (****, p<0.0001). Error bars represent standard deviation with n=3 biological replicates. (**F**) Mapping of the top mutations on the T7 RNAP structure (PDB: 3E2E). The active site is indicated by a red circle. (**G**) Heatmap showing most common T7 RNAP mutations explored by EVOLVEpro over rounds of evolution. Any position explored more than once is shown on a cumulative basis across rounds. (**H**) Scatter plot comparing the predicted ESM-2 protein fitness score versus experimentally measured T7 RNAP transcription fidelity scaled score across evolution rounds. The correlation and linear regression line are shown in the plot. (**I-J**) Comparison of the T7 RNAP latent space with either predicted ESM-2 protein fitness (masked marginal score) or EVOLVEpro protein activity fold improvement. (**K**) A kernel density estimate of protein fitness as predicted by ESM-2 versus protein activity as predicted by EVOLVEpro. The correlation and linear regression line are shown in red and the R square of correlation is reported.

During the initial two rounds of evolution, improvements were observed but were only in the 2-4 fold improvement range. However, by rounds 3 and 4, we started observing significant improvements in all features, especially in translation and immunogenicity fold changes over the wild-type T7 RNAP (Fig. 5B-C, fig. S7A). By the end of round 4, we observed one T7 RNAP mutant, E643G, that could generate luciferase mRNA that produced 34x more translated luciferase and ∼98% less immunogenicity (Fig. 5C). We sought to benchmark E643G against the previously engineered state of the art G47A/884G mutant T7 RNAP that has markedly reduced immunogenic byproduct in our IVTT assay(*47*). We found that our E643G mutant produces 7-fold higher translation in cells and ∼2-fold less IFNB1 inflammation in BJ fibroblasts (fig. S7B).

Given the plethora of mutants tested in the first 4 rounds along with the existing G47A mutation that is known to reduce dsRNA formation, we then used EVOLVEpro to generate multi-mutants that involved the combination of up to 7 previously tested mutations. Typically in rational mutagenesis for higher activity mutants, rational combinations of single beneficial mutations are combined according to their spatial location under the assumption of synergistic effects of these mutations. Here, we relied on EVOLVEpro’s ability to learn the activity landscape to nominate multi-mutants in an unbiased fashion. Surprisingly, the top nominated mutant in Round 5 is a combination of previously reported G47A and the best single mutant E643G as normal rational mutagenesis would do. However, it is worth noting that this combination resulted in a protein with worse performance than E643G alone (fig. S7B). This points to the vast unknown epistatic effect between residues at different spatial positions in a protein and the utility of EVOLVEpro in nominating multi-mutants. By round 6, EVOLVEpro was able to nominate one particular variant that had ∼57x more translation from luciferase mRNA and ∼515x less immunogenicity than the original wild-type T7 RNAP (Fig. 5C). Moreover, this variant was substantially more effective at translation and less immunogenic than the G47A/884G mutant. This multi-mutant, T7 RNAP^T3M/G47A/E643G^, was chosen as the final EVOLVEpro evolution candidate and termed enhanced T7 RNAP, or epT7.

Given the high throughput testing of mutant T7 RNAPs in the IVTT reaction, we hypothesized that the unoptimized IVT buffer could change these mutant’s mRNA production(*48*) and sought to compare the performance of top mutants in clinically relevant IVT settings with NEB’s HiScribe transcription kit, followed by Vaccinia cap-1 capping and polyA tailing. We therefore purified the top performing single mutant (E643G), previously reported state-of-the-art mutant (G47A/884G)(*47*), and our epT7 (T3M/G47A/E643G) along with WT to compare their performance. We compared the production of six different mRNA sequences, ranging in size from 500 nt to 6,500 nt, between epT7, T7^E643G^, and wild-type T7. Consistent with our IVTT-based experiment, we found that epT7 and E643G produced significantly higher mRNA in a 2-hour transcription scheme than both wild-type T7 RNAP and the G47A/884G variant (fig. S7C). Analysis of the 3 different mRNA products by both E-gel EX and TapeStation gel electrophoresis systems confirmed the presence of a single on-target product across all four enzymes (fig. S7D-F). Looking at the translation and immunogenicity aspects of the mRNAs produced by these enzymes, we found that in all cases epT7 produced mRNA had 4 to 120-fold higher translation than wild type and 4 to 256-fold lower immunogenicity (Fig. 5D, fig. S8A). Functional testing of SpCas9 mRNA also shows significantly higher editing from epT7’s produced mRNA in two separate cell lines (fig. S8B). These results validate that the EVOLVEpro derived epT7 mutants are not buffer or template-specific and are genuinely improving the quality of mRNA produced by the polymerase. We next investigated the mechanism of the epT7 performance enhancements by investigating the quality of the RNA. Using an established ELISA for dsRNA, we found that the dsRNA in the epT7-produced mRNA was 5-fold lower than wild-type T7-produced RNA and it performed equally well as the RNA produced by the state-of-the-art G47A/884G mutant(*47*) (Fig. 5E).

Previous efforts to reduce dsRNA production relied on adding a glycine residue at the C-terminal “foot” region of the enzyme (884G insertion)(*47*). Our model revealed the functional importance of E643 in transcription and, surprisingly, mutating this residue rendered the same effect as 884G (Fig. 5F, fig. S7B). Indeed, analysis of the T7 RNAP structure reveals that E643 is close to the DNA template, suggesting that E643G improves template binding and RNA production (Fig. 5F). However, E643K/E643R did not improve the fidelity of transcription (fig. S7A), suggesting that these bulky residues sterically clash with the template DNA. Therefore, it is likely that EVOLVEpro is able to identify a unique mechanism, interrogate the effect of this mechanism, and determine the right balance biochemically to mutagenize, thereby producing a novel, SOTA T7 RNAP variant that has never been described before. G47A has been previously reported to increase helix formation, and EVOLVEpro took advantage of this helix-favoring mutation in our multi-mutant generation. The third mutated residue in epT7 is in a disordered region (T3M), suggesting a role independent of DNA template binding. T3M might be involved in improving protein stability or other aspects that can modulate the polymerase activities. These residues further highlight the insightfulness of EVOLVEpro to identify novel mutants that one would not test via rational mutagenesis. Further analysis of EVOLVEpro’s residue exploration in evolution revealed that E643 was found first in round 3 with the most beneficial mutation being E643N (Fig. 5G). The model quickly gained an understanding of the functional importance of this residue and zoomed into this region by exploring it 5 more times in round 4, yielding E643G the best single mutant (Fig. 5G). This trend is similar to the evolution of proteins reported above, where a beneficial mutation at a certain residue is capitalized by the model in the next round by exploring additional mutations around that region.

We next calculated the relationship between the activity (observed data) and fitness (pMMS) for T7 RNAP and found a negative correlation of 0.13 in this case, denoting the lack of association between the two metrics. EVOLVEpro successfully navigated through this divergence by selecting mutants with higher activity but not fitness in later rounds (Fig. 5H). Lastly, we investigated the global evolutionary landscape of epT7 and EVOLVEpro’s mutational trajectory. At a high level, as with the previous proteins evolved, the activity map learned by EVOLVEpro diverged from the fitness map predicted by ESM-2, showing that fitness predictions would not be able to predict the mutants that were ultimately discovered to improve protein activity and other parameters (Fig. 5I-K).

### Circular RNA production with epT7

Circular RNA has emerged as a promising therapeutic modality for protein replacement therapy thanks to its enhanced stability and prolonged expression of proteins(*49*). Since we observed significantly lower dsRNA production and higher fidelity of transcription with epT7, we hypothesized that epT7 would enhance circular RNA production since the use of RNAse R during post-IVT processing typically enriches for both circular RNA and dsRNA species that are immunogenic (Fig. 6A). We thus applied epT7 to the circularization of four different RNA sequences, finding that the translation obtained by circRNA from epT7 is 3 to 30 fold higher than RNA produced by WT T7 RNAP (Fig. 6B, fig. S9A-D, fig. S9J). We then used TapeStation gel electrophoresis to quantify the relative ratio of circular products post IVT and found reduced long concatemer formation in the circular RNA produced by epT7 (Fig. 6C). To better understand the mechanism behind better translation of circular RNA made by epT7, we performed gel electrophoresis using 2% E-gel EX as previously validated to check for relative ratio of precursor, nicked, intermediate and full circular RNA both pre and post-RNaseR treatment (Fig. 6D). We noticed reduced intermediate and nicked byproducts in circular RNA produced by epT7, showing higher fidelity of transcription by epT7. We used the gel electrophoresis results to quantify the ratio of circular RNA across three different templates and found significantly higher circular RNA production at around 25% efficiency, which was ∼2 fold higher than the efficiency of WT T7 RNAP, higher circRNA purity, and lower concatemer production (Fig. 6E, fig. S9G-I). Lastly, we used dsRNA ELISA to detect the amount of dsRNA left in the product after RNAse R cleanup. Consistent with our hypothesis, there is a large increase in dsRNA percentage at around 1.5% from WT T7’s produced dsRNA (Fig. 6F). This dsRNA ratio is significantly reduced to 0.2% using epT7, highlighting the fidelity of epT7 during long transcription that is needed to accommodate circular RNA production (Fig. 6F). To confirm the higher stability of circular-eGFP RNA, we transfected both wild-type T7 and epT7’s produced circRNA in HEK293FT cells and imaged them 24 hours and 72 hours post-transfection (fig. S9E-F). We observed higher GFP fluorescence from epT7 than wild-type T7 RNAP and stable expression of GFP at 72 hours similar to previously reported(*49*).

**Fig. 6.**
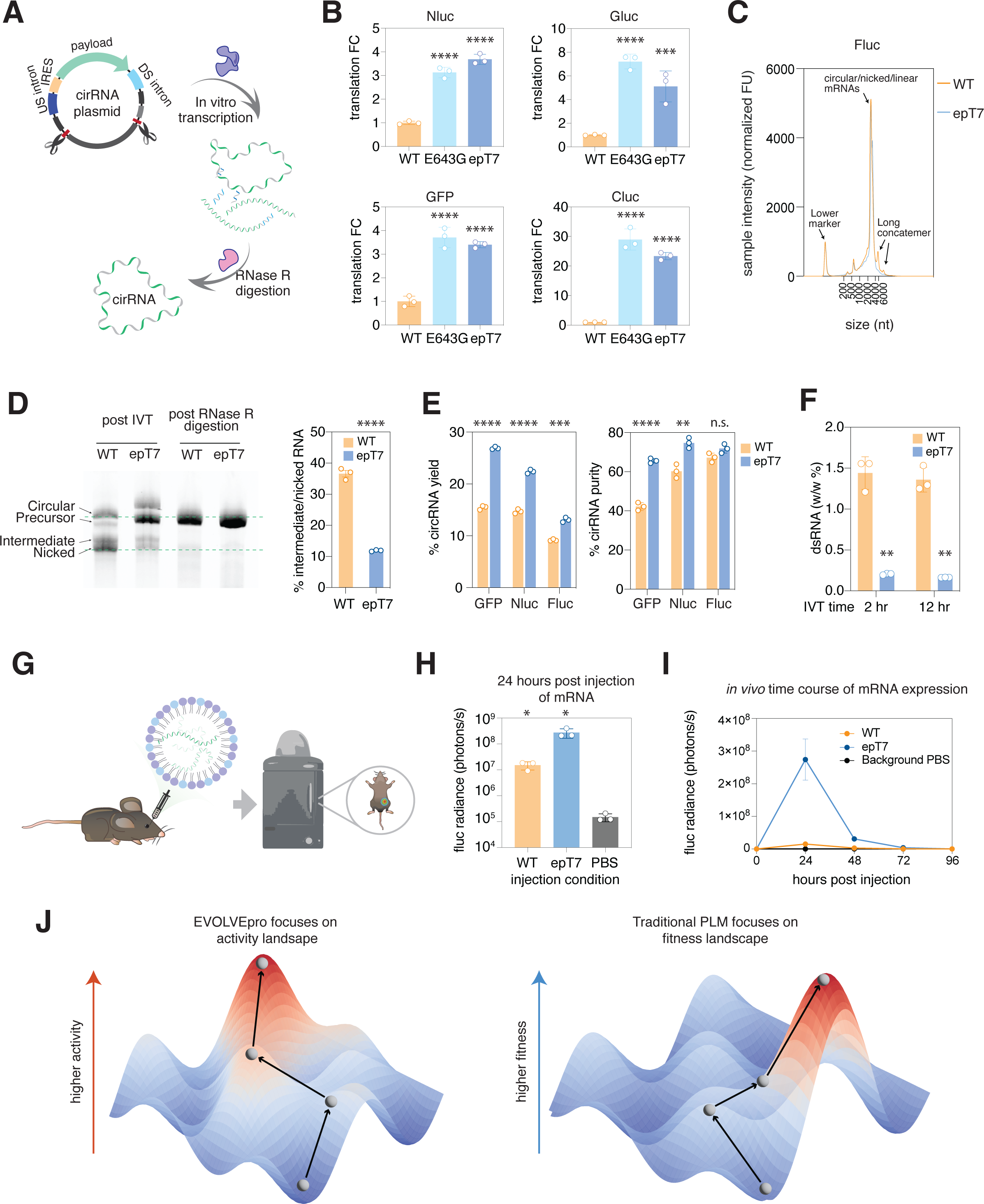
Application of epT7 for circular RNA production and *in vivo* bioluminescence. (**A**) Schematic of circular RNA production. (**B**) Validation of epT7 produced circRNA on four different template sequences compared to both T7^E643G^ and wild-type T7. Translation of each protein is measured in HEK293FT cells 48 hours after transfection. A two-sided Student’s t-test was run between WT and each evolved T7 RNAP ( ***, p<0.001, ****, p<0.0001). Error bars represent standard deviation with n=3 biological replicates. (**C**) Tapestation gel electrophoresis analysis of circular Fluc RNA produced by either epT7 or WT RNAP. epT7 shows reduced concatemer production. (**D**) Comparison of RNA products for Fluc circRNA produced by epT7 compared to wild-type T7 via gel electrophoresis using 2% E-gel EX at different steps in the production process: post-initial IVT and post-RNaseR processing. The panel on the right shows quantification of intermediate and nicked RNA ratio in the post IVT samples. Error bars represent standard deviation with n=3 biological replicates. (**E**) Comparison of purified GFP, nanoluc (Nluc), and Fluc circRNA yield by epT7 compared to wild-type T7 after the initial RNaseR clean-up. The panel on the left shows the raw mass percentage left after the cleanup. The panel on the right shows the purity of the circular RNA in the post clean-up reaction as determined by quantification using a TapeStation analysis. A two-sided Student’s t-test was run between WT and epT7 (**, p<0.01, ****, p<0.0001). (**F**) Comparison of dsRNA content for nanoluc circRNA produced by epT7 compared to wild-type T7 using either 2 hours of IVT or 12 hours of IVT. Input into the dsRNA ELISA assay involves 500 ng of post-RNAseR cleaned-up samples. A two-sided Student’s t-test was run between WT and evolved T7 RNAP (**, p<0.01). Error bars represent standard deviation with n=3 biological replicates. (**G**) Schematic of the in vivo mRNA assay for measuring mRNA expression in the liver via non-invasive luminescent imaging. (**H**) In vivo luminescent signal detected 24 hours post-injection in mice injected with mRNA produced by either epT7 or wild-type T7 or PBS controls. A two-sided Student’s t-test was run between WT, wild-type T7 RNAP, and epT7 (*, p<0.05). Error bars represent standard deviation with n=3 biological replicates. (**I**) Time-course of in vivo luminescent signal detected up to 96 hours post-injection of LNP-mRNA produced by either epT7 or wild-type T7, or PBS controls. A two-sided paired Student’s t-test was run between WT, wild-type T7 RNAP, and epT7 (*, p<0.05) for each time point. Error bars represent the standard error of mean with n=3 biological replicates. (**J**) A schematic showing the evolution of higher activity variants with EVOLVEpro. The mutagenesis landscape of proteins is often conceptualized as a complex terrain with numerous potential paths. Shown here is a gray road that conceptualizes the protein mutagenesis landscape where traversing upwards results in higher protein activity and traversing downwards reduces protein fitness. Traditional frameworks of evolutionary plausibility attempt to navigate this terrain based on natural selection, which is constrained by historical and environmental factors.

### mRNA for in vivo bioluminescent imaging

Given the high fidelity of epT7, we compared the performance of epT7 with WT T7 RNAP in producing 100% N^1^-Methylpseudouridine-5’-Triphosphate-modified firefly luciferase mRNA that is commonly used for *in vivo* deep tissue imaging (Fig. 6G). This production process, including the modified bases, mimics the clinical production of therapeutic mRNAs, allowing for a translationally relevant evaluation of epT7. We packaged the produced mRNA with lipid nanoparticles (LNPs) that traffic to the liver for bioluminescent imaging. After 24 hours post-injection of the LNP formulations, we observed ∼10-fold higher luminescence for our epT7-produced mRNA compared to mRNA produced by WT T7 RNAP (Fig. 6H). Moreover, we tracked the kinetics of both mRNA formulations for 96 hours and found consistently higher translation with the epT7-produced Fluc mRNA for a longer period of time (Fig. 6I, fig. S9K).

## Discussion

We demonstrate EVOLVEpro as an ensemble model for few-shot active learning to evolve protein activities. Over consecutive rounds of improvement, EVOLVEpro yields variants with 2- to 515-fold improvements in desired properties, including binding, catalytic efficiency, and immunogenic byproducts. Using both evolutionary scale PLMs and a regression layer, EVOLVEpro learns general rules of protein activity, generating highly active mutants with only a few cycles of evolution. Moreover, because of the rich latent space generated by PLM and powerful feature selections present in the top-layer module, EVOLVEpro evolution is a low-N learning approach that requires minimal wet lab experimentation. We benchmark EVOLVEpro across 12 different DMS datasets covering 8 protein classes, showing its superiority in the low-N evolution setting. In this benchmarking work, we evaluate all currently available embedding-based PLMs and perform a grid search to optimize over top-layer regression models, active learning selection strategies, and different normalization techniques toward the embeddings and fitness measurements. We find that PLMs are essential and their representations of protein sequence outperform traditional encoding methods like one-hot encoding and integer encoding (Fig. 1B). Interestingly, even in the extreme scarcity of data relative to the size of the input vector, dimensionality reduction of the embedding space through PCA did not improve performance (see methods and Data S1). The modular design of EVOLVEpro allows the integration of future improvements in autoregressive PLMs.

The success of EVOLVEpro speaks to the inherent limitations of PLMs, which are trained to learn a masked sequence reconstruction task across evolutionary diversity. As natural sequences do not necessarily select for optimal protein activity, the PLM’s learned fitness landscape will often not be correlated with a protein’s activity landscape. In scenarios of correlations between fitness and activity, such as antibodies, zero-shot PLM protein evolution may work with some success(*6*, *50*), but enzyme optimization has proven more challenging. It has been shown that PLMs can scale with increasing parameters just like large language models, but recent analyses have shown saturating scaling effects of PLMs with limited input training datasets (Uniref) on larger models (*51–53*). Thus, it is likely that simply increasing the parameters of these PLMs will not enable better prediction of protein activities and other downstream tasks. Alternatively, generative PLMs have yielded functional *de novo* proteins, such as GFP and CRISPR nucleases(*7*, *10*). However, variants generated by these methods do not have improved activities relative to wild-type proteins, and the functional success rate of generated proteins is low. With these limitations in mind, future generative PLMs with improved architectures and training data may be suitable for combination with EVOLVEpro to create a design framework where de novo generated sequences can be rapidly optimized for activity.

Using EVOLVEpro, we present the first comprehensive evaluation of AI protein engineering models across five therapeutically relevant proteins. These proteins have a low correlation between activity and PLM-estimated fitness, and in some cases require multiple properties to be optimized simultaneously. Critically, the assays used for measuring protein activity in this work are incompatible with pooled screening approaches, precluding typical directed evolution strategies. Across the multi-mutant landscape of protein activity, EVOLVEpro is able to select highly active single mutants out of more than 16,000 possible sequences and multi-mutants from more than 780 billion possible sequences. We thoroughly validate the five protein variants generated by EVOLVEpro for genome editing, binding, and RNA generation tasks beyond the training set, finding SOTA performance. Structural analysis of top mutations reveals many distinct mechanisms of activity improvement, suggesting future directions of improvement for these enzymes. These insights include RNaseH engineering as a means to improve prime editing and T7 polymerase engineering that potentially improves binding to the DNA template. In the context of protein design, EVOLVEpro is a highly capable protein engineering model. EVOLVEpro 1) has high rates of success, 2) requires no special knowledge about the protein, 3) can be used for multi-objective function optimization, and 4) is highly modular, allowing for any property with a quantifiable assay to be used as an input without extensive finetuning. We anticipate EVOLVEpro will continue to improve with new foundation models and enhanced search strategies and will be broadly useful for protein engineering.

## Acknowledgements

We would like to thank E. Boyden for MiSeq instrumentation support, J.Yim, A.Kirjner, I. Fiete, M. Yan, B. Li, M. Guo, T. Chen, X. Yang, C.J. Bashor, P. Mehta, and J. Rocks for computational discussions, J. Xie, G.Gao for viral preparation, Y. Shen and K. Kato for nuclease purification, R.A. Wesselhoeft for circular RNA advice, and Z. Tang, D. Irvine, J.S. Weissman, R. Desimone, and J. Crittenden for support and helpful discussions. We thank the members of the Nishimasu and Abudayyeh-Gootenberg labs for their helpful discussions.

## Authors contributions

Conceptualization: KJ, OOA, JSG

Methodology: KJ, OOA, JSG, ZY, MDB, MH, HN, BK, JKC

Investigation: KJ, OOA, JSG, ZY, AK, LV, SRS

Visualization: KJ, MDB

Funding acquisition: OOA,JSG, HN

Project administration: OOA,JSG

Supervision: OOA, JSG

Writing – original draft: KJ, OOA, JSG, ZY, MDB, MH, HN

Writing – review & editing: KJ, ZY, MDB, SRR, LV, AK, BK, JKC, MH, HN, JSG, OOA

## Data and materials availability

Expression plasmids are available from Addgene under UBMTA; support information are available within this document and supplementary materials. Models and codes are available at the following github repositories: https://github.com/mat10d/EvolvePro (full data repository) and https://github.com/idmjky/EvolvePro (deployable model)

