## Supplementary Information for "Rapid protein evolution by few-shot learning with a protein language model"

### Materials and Methods

#### Benchmarking on 12 DMS datasets

For model benchmarking, we took 9 existing deep mutational scanning (DMS) datasets which were employed in a previous zero-shot high fitness prediction approach(*1*). From this work, we leveraged a pre-determined cutoff for high-fitness variants for each dataset to select variants that were low and high fitness. To augment the use of these datasets, we also selected three additional DMS datasets: an AsCas12f compact genome editor(*2*), Cov2 viral spike receptor-binding domain(*3*), and Zika virus envelope protein(*4*). For these datasets, cutoff values were set based on the general distribution of high-activity variants. To facilitate downstream work, a tabular format CSV file and a fasta file of all available mutant sequences with activity measurements were generated from each dataset.

#### Extraction of PLM embeddings

From the set of all available protein variants in the 12 DMS datasets, protein language model embeddings were generated. We optimized the use of ESM(1–4), ProtTrans(5), ProteinBERT(6), Ankh(*5*), and UniRep(*6*) methodologies to operate on fasta files of all available mutants. These were used with default parameters. In the case of ESM embeddings, mean embeddings were generated, for esm1b, esm1v, and esm2 models of varying size. For ProtTrans, a per-protein embedding was generated. For ProteinBERT, global representations were generated. For Ankh, the mean of the last hidden state was taken for both the base and large models. For UniRep, the h_avg was generated. In all of these cases, a vector-based representation of each protein variant was the final objective. In addition to this, one-hot encodings and integer encodings of all variants were generated. In order to preserve the single n-dimensional vector embedding for each protein, one-hot encodings were concatenated across all positions.

#### EVOLVEpro Parameter Grid search

We conducted an extensive grid search to evaluate various strategies for optimizing fitness in a low number of rounds. The grid search explored the following parameters:

1. Fitness measurement: Raw fitness values from each dataset or min-max normalized fitness.
2. First-round strategy: Random selection of variants or diverse selection using K-medoids clustering on protein language model embeddings.
3. Learning strategies: We compared several strategies for selecting variants in subsequent rounds, including:
    a. Random selection
    b. Top n predicted fitness variants
    c. Top n/2 and bottom n/2 predicted fitness variants
    d. Maximizing Euclidean embedding distance from previously selected variants
4. Embedding types: We compared different embedding representations, including raw embeddings, normalized embeddings across residues, and PCA-reduced (10 PCs) embeddings to account for the fact that this was a high p, low n paradigm. This was entirely done on the largest (15B parameter) ESM2 model.
5. Regression types: We evaluated various regression models for fitness prediction, including ridge regression, lasso regression, elastic net, linear regression, neural networks with a linear last layer, random forest regression, and gradient boosting regression. These were largely used with default parameters.

For each combination of parameters, we ran three simulations (to vary the first round of selected variants) using 16 variants per round to account for stochastic variability. Performance was assessed using the proportion of high-fitness variants out of the top 16 variants that the updated model would predict. We quantified the overall effectiveness of each parameter value by counting the number of datasets for which it achieved the highest mean fitness binary percentage. This "winning strategy" count provided a simple yet informative summary of which approaches were most successful across diverse protein systems. The “winning strategy” was: random first round, raw fitness, top-n selection, random forest regression, and raw embeddings.
PLM comparison:
Lastly, to understand if the largest ESM2 (15B parameter) model was optimal for our EVOLVEpro model, we sought to understand if our “winning strategy” would be aided by using a different underlying PLM. For each PLM, we ran ten simulations of our winning strategy and assessed this using the proportion of high-fitness variants out of the top 16 variants that the updated model would predict over 10 rounds.

EVOLVEpro Model

In EVOLVEpro, we utilize a Random Forest regressor as our top layer model to learn the functional grammar of proteins based on information-rich latent space embeddings generated by a protein language model in an active learning setting. Let $x_i$ denote the amino acid sequence of the i-th protein variant. The protein language model embedding transformation transforms $x_i$ into a d dimensional vector representation:

$$E(x_i) = [e_1, e_2, \ldots, e_d]^d$$

To reduce the number of features fed into the random forest regressor in the low-N setting, we obtain the average embedding vector for all variants by computing the mean of the embeddings across all residues for a given protein at length n:

$$[\bar{E}(x_i) = \frac{1}{n_i} \sum_{j=1}^{n_i} e_{ij}]$$

The Random Forest model f then operates on these reduced embeddings to predict the fitness value $y_i$ of each protein variant. Each decision tree $h_t$ within the Random Forest makes a prediction based on recursive binary splits of the input latent space features based on a defined threshold ϴ and the composite function average across T individual decision trees.

$$f(\bar{E}(x_i)) = \frac{1}{T} \sum_{t=1}^{T} h_t(\bar{E}(x_i))$$

$$h_t(\bar{E}(x_i)) = \begin{cases} c_L & \text{if } \bar{E}(x_i)_j \leq \theta \\ c_R & \text{otherwise} \end{cases}$$

Then this top layer domain expert model is trained to minimize the Mean Squared Error (MSE) between the predicted and actual activity values in a round-to-round active learning fashion where N varies from 10 to 100:

$$MSE = \frac{1}{N}\sum_{i=1}^{N}(y_i - f(\bar{E}(x_i)))^2$$
$$y_{pred} = f(\bar{E}(x_{new}))$$

Comparison with zero-shots model

To assess whether our model would perform better than other one-shot strategies, we also compared 5 and 10 rounds of evolution through EVOLVEpro to a one-shot round of pre-training on 16, 80, 160, 500, and 1000 mutants. In this approach, we essentially conducted a single random round of this size, and again assessed using the proportion of high-fitness variants out of 16 variants predicted in this one-shot format, across 10 simulations of the initial round. We also compared a default implementation of a zero-shot high fitness prediction approach(*1*), where we generated variants for the 3 additional datasets not assessed in this paper for full benchmarking. Given that this one-shot approach was quite narrow, in selecting less than 32 variants, and that ultimately across 5 rounds of our strategy, we were pulling 80 variants, we manipulated parameters in this zero-shot approach to attempt to select more than 80 variants for better benchmarking.

Comparison of few shots versus many shots

We logically assumed that EVOLVEpro would improve if more rounds and more variants p er round were provided to the underlying grid searches, given that with more training data our m odel would improve. Therefore, we did not vary these parameters in the initial grid search. Howe ver, to observe how the varying round size would affect the saturation of selecting only high-fitne ss variants, we tested 10, 20, 30, 40, 50, 100, 200, and 500 variants per round using the "winning strategy" across all 12 datasets. Given the size of certain DMS datasets, this was capped at 100 an d 200 variants per round for certain datasets for which no more than 1000 or 2000 variants were a vailable. All in-silico optimization of EVOLVEpro is available at EVOLVEpro

Experimental use of EVOLVEpro

Given the optimization of EVOLVEpro done across the 12 DMS datasets, we sought to apply our "winning strategy" across a range of protein fitness and activity optimization tasks with experimental output. In this work, the "minimal strategy" was put into use, with the caveat that raw fitness was not usable given the round-by-round experimental fitness variation. For PsaCas12f evolution, ESM1b-650M base layer PLM was used for evolution. To ensure normalization across rounds, fitness was normalized to wild-type fitness in each round. Furthermore, for T7 RNAP evolution, we attempted two multi-mutant rounds, where we generated variants with multiple amino acid substitutions that included all single amino acid substitutions that had a fitness greater than wild-type fitness. During the experimental setup, the top 10 to top 12 mutants were selected for testing to better suit the high throughput setup. The model for experimental evolution is available at https://github.com/idmjky/EVOLVE-Pro

ESM2 fitness calculation

We use the previously established EMS2' masked marginal scoring function(*7*):

$$\sum_{i \in M} \log p(x_i = x_i^{\text{mt}} \mid x_{-M}) - \log p(x_i = x_i^{\text{wt}} \mid x_{-M})$$

where M are the masked residues where mutations occur, $x_i^{mut}$ is the mutant-type residue at position i, and $x_i^{WT}$ is the wild-type residue at position i. This function was shown to perform best. Embeddings are computed with ESM2-15B and then the individual mutant's fitness score is calculated by the function above. The code for performing this calculation is available within the GitHub repositories.

Cas12f protein purification and preparation

The gene encoding PsaCas12f (residues 1–586) with an N-terminal $His_6$-SUMO tag was cloned into the pE-SUMO vector (LifeSensors). The mutations were introduced by a PCR-based method, and the sequences were confirmed by DNA sequencing. The Cas12f protein was expressed in *Escherichia coli* Rosetta2 (DE3) (Novagen) by induction with 0.25 mM isopropyl β-D-thiogalactopyranoside (Nacalai Tesque) at 20°C overnight. The E. coli cells were lysed by sonication in buffer A (20 mM Tris-HCl, pH 8.0, 20 mM imidazole, 1 M NaCl, 3 mM 2-mercaptoethanol, 10% glycerol) and the lysate was clarified by centrifugation at 40,000 × g. The supernatant was applied to Ni-NTA Superflow resin (QIAGEN), and the Cas12f protein was eluted with buffer B (20 mM Tris-HCl, pH 8.0, 300 mM imidazole, 300 mM NaCl, 3 mM 2-mercaptoethanol, 10% glycerol). The eluate was treated with Ulp1 peptidase at 4°C overnight and then loaded onto a HiTrap SP column (GE Healthcare), equilibrated with buffer C (20 mM HEPES, pH 7.5, 300 mM NaCl, 2 mM DTT). The protein was eluted with a linear gradient of 0.3–2 M NaCl. The Cas12f protein was further purified on a Superdex 200 Increase 10/300 column (GE Healthcare), equilibrated with buffer D (20 mM HEPES, pH 7.5, 1 M NaCl, 2 mM DTT). The peak fractions were collected and stored at -80°C until use. The sgRNAs were transcribed in vitro with T7 RNA polymerase, using PCR-amplified DNA templates, and were purified by 10% denaturing (7 M urea) polyacrylamide gel electrophoresis.

*In vitro* DNA cleavage assay

The purified PsaCas12f was diluted to 2 µM (20 mM HEPES, pH 7.5, 600 mM NaCl, 2 mM DTT) and mixed with an equal volume of the sgRNA (2 µM) at 37°C for 2 min. The pre-assembled PsaCas12f–sgRNA complex (5 µL, 1 µM) was then mixed with the linearized plasmid target containing the target sequence and the TTA PAM (2.5 µl, 100 ng/µL) and buffer F (2.5 µL, 60 mM HEPES, pH 7.5, 40 mM $MgCl_2$, 2 mM DTT). The 10 µL reaction solution (500 nM PsaCas12f–sgRNA, 250 ng target DNA, 20 mM HEPES, pH 7.5, 150 mM NaCl, 10 mM $MgCl_2$, 1 mM DTT) was incubated at 37°C. Aliquots (2 µL) were taken at 15, 30, and 60 min, and mixed with 6 µL of quench solution (20 mM HEPES, pH 7.5, 150 mM NaCl, 2 mM DTT, 3.5 µg Proteinase K, 17.5 mM EDTA). The reaction products were incubated at 95°C for 2 min and then analyzed using a MultiNA microchip electrophoresis system (SHIMADZU).

Measurement of luciferase activity

Media containing secreted or intracellular luciferase was harvested 48 hours after transfection unless otherwise noted. 20µL of media is used to measure secreted luciferase activity using Targeting Systems Cypridinia and Targeting systems Gaussia luciferase assay kits (Targeting Systems) on a Biotek Synergy 4 plate reader with an injection protocol. All replicates were performed as biological replicates. Intracellular Nanoluc and firefly luciferase were measured by lysing the cell in the luciferase assay mix (Promega) according to the manufacturer's protocol. 5 minutes after lysis at room temperature, the signal is read out using a Biotek Synergy Neo2 plate reader.

Quantification of protein expression

Two days after the transfection of HEK293FT or BJ Fibroblast cells, the Nano-Glo HiBiT Lytic Detection System (Promega) was used for the quantification of the HiBiT tags, in cell lysates. For the preparation of the Nano-Glo HiBiT Lytic Reagent, the Nano-Glo HiBit Lytic Buffer (Promega) was mixed with Nano-Glo HiBiT Lytic Substrate (Promega) and the LgBiT Protein (Promega) according to the manufacturer's protocol. The volume of Nano-Glo HiBiT Lytic Reagent added was equal to the culture medium present in each well, and the samples were placed on an orbital shaker at 600 rpm for 3 minutes. After incubation of 10 minutes at room temperature, the readout took place with 125 gain and 2 seconds integration time using a plate reader (Biotek Synergy Neo2). The control background was subtracted from the final measurements.

Harvest of total RNA and quantitative PCR

For gene expression experiments in mammalian cells, cell harvesting and reverse transcription for cDNA generation were performed using a previously described modification of the commercial Cells-to-Ct kit (Thermo Fisher Scientific) 48 h after transfection. Transcript expression was then quantified with qPCR using Fast Advanced Master Mix (Thermo Fisher Scientific) and TaqMan qPCR probes (Thermo Fisher Scientific) with GAPDH control probes (Thermo Fisher Scientific). All qPCR reactions were performed in 10-µl reactions with two technical replicates in a 384-well format and read out using a LightCycler 480 Instrument II (Roche). For multiplexed targeting reactions, readout of different targets was performed in separate wells. Expression levels were calculated by subtracting housekeeping control (GAPDH) cycle threshold (Ct) values from target Ct values to normalize for total input, resulting in ΔCt levels. Relative transcript abundance was computed as 2−ΔCt. All replicates were performed as biological replicates.

AAV production and purification

Recombinant AAV2/8 was produced by transient HEK293 cell transfection and CsCl sedimentation by the University of Massachusetts Medical School Viral Vector Core, as previously described(*8*). Vector preparations were monitored by ddPCR, and purity was assessed by 4%–12% SDS-acrylamide gel electrophoresis and silver staining (Invitrogen).

Animal AAV injection and processing

For *in vivo* testing of enPsaCas12, an AAV was prepared at a titer of $1.5 \times 10^{13}$ gc/mL in sterile phosphate-buffered saline PBS (University of Massachusetts Medical School Viral Vector Core). Animals were randomly assigned to experimental or control groups. The investigator was not blinded to assignments. AAV was delivered to 3-month-old male C57/BL6 mice via retro-orbital injection at a dose of $1.5 \times 10^{12}$ genomic copies adjusted to 100 µL with PBS, pH 7.4 (Gibco), before the injection. In control animals, 100 µL of sterile PBS was administered via retro-orbital injection. To monitor serum levels of PCSK9 and total cholesterol, blood was routinely drawn from mice pre- and post-injection. Mice were fasted for 12 hours overnight prior to the blood draw. Blood collection was performed by saphenous vein sampling, with no more than 1% of the blood volume collected over a 24-hours period. To collect serum, whole blood was incubated at room temperature to allow clotting for 1 h, followed by centrifugation at 10,000 × g for 10 min. Serum samples were used immediately for testing, with the remaining samples stored at −20°C for any subsequent analysis. Animal studies were performed in accordance with the recommendations in the Guide for the Care and Use of Laboratory Animals of the National Institutes of Health. The protocols were approved by the Institutional Animal Care and Use Committee at the Massachusetts Institute of Technology.

*In vivo* Firefly luciferase mRNA delivery and comparative in vivo bioluminescence

Prior to bioluminescence imaging, 8 to 10-week-old Albino B6 were anesthetized with 3% isoflurane and injected with 5 μg of synthesized mRNA via retro-orbital injection using homemade lipid nanoparticles. At the indicated time points post-injection, the mice were anesthetized again with 3% isoflurane and immediately administered 200 µl of 15 mg/mL D-luciferin (PerkinElmer) for imaging. Ventral bioluminescence images were acquired using an IVIS Spectrum In Vivo Imaging System (PerkinElmer). The following conditions were used for image acquisition: exposure time = 60 sec, binning = medium: 4, field of view = 15 x 15 cm, and f/stop = 1. Bioluminescent images were analyzed using Living Image 4.3 software (PerkinElmer) and normalized radiance (photons/s) was reported.

Serum and tissue analysis

To quantify serum PCSK9, the Mouse Proprotein Convertase 9/PCSK9 Quantikine ELISA Kit (R&D Systems) was used with fresh serum samples, according to the manufacturer's protocol. Total cholesterol levels were measured with the Cholesterol/Cholesterol Ester-Glo™ Assay (Promega), according to the manufacturer's protocol. All assays were performed using a Biotek Synergy Neo2 plate reader. To assess genome editing in livers, mice were euthanized by carbon dioxide inhalation. Livers were extracted and placed in ice-cold, sterile PBS. The liver portions that were not used for immediate analysis were snap-frozen and stored at −80°C. To isolate genomic DNA, liver pieces were processed using the DNeasy Blood & Tissue Kit (Qiagen), according to the manufacturer's protocols. The *PCSK9* region of interest was amplified from purified genomic DNA and sequenced as described above.

T7 RNA polymerase purification

Plasmids for overexpressing Twin-Strep-tagged SUMO (Small Ubiquitin-like Modifier)-fused WT or mutant T7 polymerase (pET-6xHis-thrombin-Twin-Strep-tag-SUMO-T7) were transformed into BL21 T7 expression *E. coli* strain (NEB, C2566H). The transformed cells were inoculated into 1.2 liters of Terrific Broth (TB) with 100 μg/ml ampicillin using 12 ml of an overnight culture of T7 Express cells containing the T7 polymerase expression construct. Cultures were grown at 37 °C until the cell density reached OD600 ~0.6, then protein overexpression was induced by adding 0.2 mM isopropyl β-D-thiogalactoside (IPTG) and incubating for 24 hours at 16 °C. Cells were harvested by centrifugation at 4,000 g for 15 minutes and stored at -80 °C until purification.

The cell pellet was resuspended in lysis buffer (50 mM Tris-HCl, pH 8.0, 500 mM NaCl, 1 mM DTT) containing Protease Inhibitor (Roche Complete ULTRA, EDTA-free), lysozyme (Thermo Fisher Scientific), and Benzonase (Millipore) to degrade nucleic acids after lysis. Cells were lysed using an ultrasonic homogenizer under ice-cooling, followed by clarification through 60 minutes of centrifugation at 10,000 g. The lysate was incubated with Strep-Tactin®XT resins (iba) at 4 °C for 1 hour with orbital shaking, then applied to a gravity flow chromatography column equilibrated with wash buffer (50 mM Tris-HCl, pH 8.0, 500 mM NaCl, 1 mM DTT, Protease Inhibitor). After washing, the protein was eluted with the same cleavage buffer (50 mM Tris-HCl, pH 8.0, 500 mM NaCl, 0.1% Triton® X-100, 1 mM DTT, SUMO protease) following overnight cleavage of the SUMO-fusion by SUMO protease (Sigma-Aldrich) at 4 °C. Proteins were concentrated to 500 ml using Amicon® Ultra Centrifugal Filters (30 kDa molecular-mass cut-off, Millipore) before loading onto a gel filtration column (Superdex® 200 Increase 10/300 GL) via FPLC (AKTA Pure). Gel filtration fractions were analyzed by SDS-PAGE, and those containing T7 polymerase were pooled.

Proteins were quantified using the CBQCA Protein Quantitation Kit (Thermo Fisher Scientific) per the manufacturer's instructions, buffer exchanged into storage buffer (50 mM Tris-HCl, pH 8.0, 100 mM NaCl, 1 mM DTT, 0.1 mM EDTA, 50% Glycerol, 0.1% Triton® X-100), and stored at -20 °C until use.

Linear mRNA production

IVT templates for linear mRNA synthesis were prepared either by PCR amplification for 35 cycles or by linearizing plasmid DNA with PmeI (NEB, R0560L) overnight. The resulting products were purified using silica columns (Qiagen, 28006 for PCR products or 28115 for plasmid digestion products) before RNA synthesis. Linear mRNA was synthesized *in vitro* at 37°C for 2 hours using 1 μM purified T7 polymerase with either 1000 ng of linearized plasmid template or 500 ng of PCR-amplified DNA templates per 20 µL of IVT reaction. Equimolar concentrations of NTPs (NEB, N0466L) were used, and the reactions for wild-type and mutant T7 polymerase were performed under identical conditions using 10x T7 buffer (NEB, E2040). For *in vivo* tests, mRNA was synthesized in vitro by T7 RNAP-mediated transcription at 37°C for 4 hours using 100% substituted N1-methylpseudouridine-triphosphate (TriLink, N-1019) with either 1 μM purified epT7 polymerase or 2 µL of commercially available WT T7 RNA Polymerase (NEB, M0251S) per 20 µL using the 10x T7 buffer from NEB HiScribe transcription kit (NEB, E2040). DNase I (NEB, M0303L) was used to remove the DNA template, terminating transcription. The mRNA was then purified using a silica column (NEB, T2040L) and quantified using a Nanodrop One Microvolume UV spectrophotometer (Thermo Fisher, ND-ONE-W) before further enzymatic reactions. The Cap 1 structure was added to the 5′ end using Vaccinia capping enzyme (NEB, M2080) and mRNA Cap 2'-O-Methyltransferase (NEB, M0366). The mRNA was purified again using a silica column (NEB, T2040L), followed by the addition of AMP from ATP to the 3′ end using E. coli Poly(A) Polymerase (NEB, M0276L). All mRNAs were column purified (NEB, T2040L) and eluted with 1 mM sodium citrate (Thermo Fisher, AM7001).

Circular RNA production

The construction of the plasmid template for circular mRNA synthesis has been previously described(*9*). The plasmid was digested with NotI restriction enzyme (Thermo Fisher, FD0593), and the resulting DNA product, serving as the transcriptional template for circular RNA, was column purified using a MinElute PCR Purification Kit (Qiagen, 28006). For each 20 µl IVT reaction, 500 ng of the purified transcriptional template was used. The IVT reactions were incubated at 37°C for 12 hours using 10x T7 buffer (NEB, E2040), followed by degradation of the DNA template with 2 µl of DNase I per 500 ng of transcriptional template for 30 minutes at 37°C. For epT7, 2 μM of the purified protein is used as input and for WT, 2 µL of the wild-type T7 RNA polymerase (NEB, M0251S) is used as input. The remaining RNA was column purified before further enzymatic processing.

A separate circularization step was performed to promote covalently closed circular RNA formation from the uncircularized IVT product. This step included 1X T4 RNA ligase I buffer (NEB, B0216L), 2 mM GTP (NEB, N0450L), and 20 units of RNase inhibitor (NEB, M0314L) incorporated into 45 µg of silica column-purified RNA in a 50 µl reaction. The reaction mixture was heated at 55°C for 8 minutes and then silica column purified (NEB, T2040L). To isolate circular RNAs, the column-purified RNA was digested with 4 units of RNase R (Abcam, ab286929) per microgram of RNA for 15 minutes at 37°C. The samples were subsequently column purified, quantified using a Nanodrop One Microvolume UV spectrophotometer, and verified for complete digestion using the E-Gel Electrophoresis System or an Agilent TapeStation, following the manufacturer's instructions.

Antibody and antigen production

For high throughput antibody production, we transfected HEK293FT cells with 50 ng of heavy chain encoding plasmid and 50 ng of light chain encoding plasmid 24 hours after seeding at 20,000 cells per well in a 96-well plate. Antibodies are harvested 72 hours after transfection by centrifuging at 4,000x g for 10 minutes and supernatants were directly used for downstream ELISA. Here, we chose the previously established REGN10987 heavy chain only for mutation(*1*).

For antigen purification, SP6 stabilized antigen was His-tagged and purified using HisPur Ni-NTA resin (Thermo Fisher Scientific, 88222). Cell supernatants were diluted with 1/3 volume of wash buffer (20 mM imidazole, 20 mM 4-(2-hydroxyethyl)-1-piperazineethanesulfonic acid (HEPES) pH 7.4, 150 mM sodium chloride (NaCl) or 20 mM imidazole, 1× PBS), and the Ni-NTA resin was added to diluted cell supernatants. Resin/supernatant mixtures were added to chromatography columns for gravity flow purification. The resin in the column was washed with wash buffer (20 mM imidazole, 20 mM HEPES pH 7.4, 150 mM NaCl or 20 mM imidazole, 1× PBS), and the proteins were eluted with 250 mM imidazole, 20 mM HEPES pH 7.4, 150 mM NaCl or 20 mM imidazole, 1× PBS. Column elutions were concentrated using centrifugal concentrators at 10-kDa, 50-kDa, or 100-kDa cutoffs, and stored in a storage buffer at -20°C.

High throughput Antibody concentration determination and ELISA

We adopted a previously developed high-throughput ELISA where we coated protein binding titer plates with 50uL of polyclonal goat-anti-human IgG at 2.5 µg/ml in 1× PBS overnight at 4°C(*10*). After overnight coating, we removed the coating solution by flicking the plate and tapping the residual liquid on paper towels. We washed each well of the ELISA plate six times with 200 µl of washing buffer and removed the remaining fluids by flicking the plate and blocking the plate with 200 µl of blocking buffer per well for at least 45 min at 37 °C. We then washed each well of the ELISA plate six times with 200 µl of washing buffer and flicked the remaining fluids. A standard human myeloma IgG1 kappa at 4 µg/ml in 1× PBS was used to create a serial dilution of standards and added in with dilutions of produced antibodies for 45 minutes at room temperature. We wash each well of the ELISA plate six times with 200 µl of washing buffer and flick the remaining fluids. We then diluted HRP-conjugated secondary antibody 1:1,000 in a blocking buffer and added 50uL of the secondary antibodies to each well. Plates were incubated at room temperature for 45 minutes. After incubation, plates were washed again six times with wash buffer, and 50µl of ABTS solution at room temperature was added per well. 5 minutes after addition, 2M sulfuric acid is added to stop the reaction and OD450 is read out using a Biotek plate reader. For the binding activity of the antibody, the same protocol is used except coating the plate with 100nM of purified antigen at 4°C overnight.

dsRNA ELISA

The dsRNA byproduct from *in vitro* transcription was detected using a dsRNA sandwich enzyme-linked immunosorbent assay (ELISA) that selectively identifies multi-species dsRNA molecules larger than 30-40 bp. This assay was performed with a multi-species dsRNA ELISA Kit (Novus Biologicals, NBP3-11368), using antibodies as previously described by Schonborn, J., et al. The K1 (IgG2a) mouse monoclonal antibody was immobilized on 96-well Immulon 2 HB plates (Thermo Fisher Scientific, 3455) overnight at 4°C, then blocked with 1% BSA in PBS at 37°C for 2 hours. After triple washing, mRNA samples and Poly (I:C) dsRNA standards were added to the plates and incubated for 1 hour at 37°C. Following another triple wash, the plates were incubated with monoclonal antibody K2 (IgM) at 37°C for 1 hour. After triple washing again, the plates were exposed to horseradish peroxidase (HRP)-conjugated F(ab')2 fragment of goat anti-mouse secondary antibody at 37°C for 1 hour, followed by a final wash. Subsequently, 100 µL of TMB (3,3',5,5' tetramethylbenzidine) substrate solution was added to each well, followed by 100 µL of 2M $H_2SO_4$. The absorbance was measured at 450 nm using a BioTek Synergy Neo2 Hybrid Multimode Reader (BioTek, BTNEO2).

mRNA analysis by E-gel EX system and Tapestation

Linear RNA or isolated circular RNA was column purified and quantified using a NanoDrop One spectrophotometer. For RNA characterization via the E-gel system, 1200 ng of RNA samples and 4 µL ssRNA Ladder (NEB, N0362S) were denatured by a 1:1 dilution with formamide (Sigma, F7503-250ML). The samples were then loaded onto 2% E-Gel™ EX Agarose Gels with SYBR-GOLD II and run on the E-Gel Power Snap Plus Electrophoresis System (Thermo Fisher, G9101) using the settings: E-gel Category "11 wells," E-gel Type "E-Gel™ EX 2%," and Time "12 minutes" at room temperature. Images were captured using a Bio-Rad ChemiDoc Imaging System with the "SYBR-Gold" settings. For the characterization of circular RNA using the Agilent TapeStation, 2 µL of 10 ng/µL mRNA samples and RNA ladder were mixed with 1 µL of High Sensitivity RNA Sample Buffer. The samples were denatured at 72°C for 3 minutes and then cooled at 4°C for 2 minutes. They were subsequently loaded into the 4200 TapeStation instrument (Agilent, G2991BA) and analyzed using the 4200 TapeStation Controller Software, following the manufacturer's instructions.

IVTT for T7 RNA polymerase production

For high throughput T7 RNAP generation, we used the TnT Quick Coupled Transcription/Translation System (Promega #2080) by cloning mutant T7 RNAP under an SP6 promoter to use for in vitro coupled transcription-translation. We incubated the input plasmid with the reaction mixture for 90 minutes according to the manufacturer's protocol and subsequently used 2 µL of the reaction mixture as the input T7 RNA polymerase for in vitro transcription of downstream mRNA. For IVT, we used the NEB ARCA HiScribe co-transcriptional kit with the manufacturer-supplied Cluc mRNA template. We followed the protocol by incubating the reaction mixture for 30 minutes at 37 °C. After IVT, 2 µL of DNase I was added to degrade the DNA template, and polyA polymerase was added for polyA tailing at 37 °C for 30 minutes. mRNA is purified using monarch mRNA cleanup column (#T2047, NEB) according to the manufacturer's protocol and eluted in mRNA storage buffer (#AM7000, ThermoFisher). mRNA is used directly for transfection using messengerMax (#LMRNA001, ThermoFisher), or frozen at -80 °C for subsequent characterization.

Lipid nanoparticles production

LNPs were prepared using a vortex mixing method(*11*). ALC-0315, DSPC, cholesterol, and DMG-PEG-2000 were dissolved in ethanol at 75, 10, 10, and 10 mg/ml, respectively, and mixed at a molar ratio of 50:10:38.5:1.5. The final volume was brought to 30 µl with ethanol. Separately, 10 µg of purified mRNA was diluted in 10 mM citrate buffer (pH 4) to a final volume of 90 µl. The lipid mixture (30 µl) was rapidly pipetted into the vertexing RNA solution (90 µl) at a 1:3 volume ratio, and vortexed for an additional 20 seconds. The mixture was left at room temperature for 10-15 minutes, then dialyzed against PBS at 4°C overnight to remove ethanol and residual lipids, resulting in the final LNP formulation. RNA encapsulation efficiency was measured using the Thermo Fisher RiboGreen assay. LNP samples were treated with Tris-EDTA or Tris-EDTA + 1% Triton-X. Free RNA was detected using an RNA-binding fluorescent dye, RiboGreen, and fluorescence was measured with a plate reader. Encapsulation efficiency was quantified by comparing the fluorescence of treated samples to RNA standards.

Mammalian cell culture and transfection

Mammalian cell culture experiments were performed in the HEK293FT (Thermo Fisher Scientific), BJ Fibroblast (ATCC CRL-2522), and Hepa 1-6 (ATCC CRL-1830) cell lines, grown in Dulbecco's Modified Eagle Medium with high glucose, sodium pyruvate, and GlutaMAX (Thermo Fisher Scientific), and supplemented with 1 × penicillin-streptomycin (Thermo Fisher Scientific) and 10% fetal bovine serum (VWR Seradigm). All cells were maintained at confluency below 80%. All transfections were performed with Lipofectamine 3000 (Thermo Fisher Scientific). Cells were plated 16–20 hours prior to transfection to ensure 90% confluency at the time of transfection. For 96-well plates, cells were plated at $2\times10^4$ cells/well. For each well on the plate, transfection plasmids were combined with OptiMEM I Reduced Serum Medium (Thermo Fisher Scientific) to a final volume of 10 µL.

Mammalian genome editing

To measure genome editing activity, 50 ng of protein expression construct, 50 ng of the corresponding guide construct, and optionally 20 ng of luciferase reporter were transfected in one well of a 96-well plate, using Lipofectamine 3000. After 72 h, the cells were washed once with 1 × DPBS (Sigma Aldrich). The cells were resuspended in 50 µL QuickExtract DNA Extraction Solution (Lucigen) and cycled at 65°C for 15 min, 68°C for 15 min, and then at 95°C for 10 min for lysis. A 2.5 µL aliquot of the cell lysate was used as input for each PCR reaction. For library amplification, target reporter regions were amplified with a 12-cycle PCR using NEBNext High Fidelity 2 × PCR Master Mix (NEB) with an annealing temperature of 63°C for 15 s, followed by a second 20-cycle round of PCR to add Illumina adapters and barcodes. The libraries were gel extracted and subject to paired-end sequencing on an Illumina MiSeq with Read 1 220 cycles, Index 1 8 cycles, Index 2 8 cycles, and Read 2 80 cycles. Insertion/deletion (indel) frequency was analyzed using CRISPResso2(*12*). All guide sequences are available in Data S5.

Three Primer NGS for PASTE

To quantify the integration of *attB/attP* pairs in the Bxb1 assay and genome editing for prime editing and HDR integration in the genomic locus, target regions were PCR amplified and analyzed by deep sequencing. Genomic DNA samples were isolated using 50 µl of QuickExtract (Lucigen) per well, and target regions were PCR amplified with NEBNext High-Fidelity 2× PCR master mix

(NEB) based on the manufacturer’s protocol. Barcodes and adapters for Illumina sequencing were added in a subsequent PCR amplification. Amplicons were pooled and prepared for sequencing on a MiSeq (Illumina). Reads were demultiplexed and analyzed with appropriate pipelines. To analyze the Bxb1 integration assay, the frequency of recombined *attB/attP* sites was counted relative to intact *attB* sites. To analyze prime and HDR editing, amplicons were analyzed using CRISPresso2 to count the relative number of reads with the inserted sequence. We also developed a three-primer NGS assay to quantify left junction integration using a common forward primer, a reverse primer to detect the unintegrated genomic locus, and another reverse primer for detecting the insertion template. This assay was performed as above with each reverse primer at half concentration. All NGS primers are available in Data S6.

<u>Cloning of twinPE pegRNA</u>

pegRNA were cloned by Golden Gate assembly of PCR products. Guide products were amplified by PCR (KAPA HiFi HotStart DNA polymerase, Roche) off of the Cas9 single guide RNA scaffold, with the forward primer containing spacer sequences and the reverse primer containing desired PBS, RT, and *attB* insertion sequences, in the case of the pegRNA. PCR products were purified by gel extraction (Monarch gel extraction kit, NEB) and assembled in a Golden Gate assembly containing 6.25 ng of pU6-atgRNA-GG-acceptor (Addgene, 132777), purified PCR product (approximately two- to four fold molar excess), 0.125 µl of Fermentas Eco31I (Thermo Fisher Scientific), 0.0625 µl of T7 DNA ligase (Enzymatics), 0.0625 µl of 20 mg ml–1 bovine serum albumin (NEB), 2× reaction ligation buffer (Enzymatics) and water, for a 6.25-µl total reaction volume. Reactions were incubated between 37 °C and 20 °C for 5 min each for a total of 15 cycles. Two microliters of assembled reactions were transformed into 20 µl of competent Stbl3 cells generated by Mix and Go! competency kit (Zymo) and plated on agar plates supplemented with appropriate antibiotics. After overnight growth at 37 °C, colonies were picked into Terrific Broth (TB) medium (Thermo Fisher Scientific) and incubated with shaking at 37 °C for 24 h. Cultures were collected using a QIAprep Spin Miniprep kit (Qiagen) according to the manufacturer’s instructions.

<u>High throughput Cloning of mutants</u>

Expression constructs for antibody, Bxb1 integrase, PsaCas12f nucleases, prime editor, and T7 RNAP were cloned for mammalian expression via Gibson cloning using Hifi Assembly mix (NEB) according to the manufacturer’s instructions. Overlapping reverse and forward primer-carrying mutations for desired amino acids are used to amplify the plasmid around the globe with 18bp of homology. Then DPNI is used to clean up the plasmid from PCR reactions followed by column cleanup. 50ng of the cleaned-up PCR product is then used to perform Gibson reactions according to the manufacture’s protocol. For all Gibson cloning, 2 µl of assembled reactions were transformed into 20 µl of competent Stbl3 cells generated by Mix and Go! competency kit (Zymo) and plated on agar plates supplemented with appropriate antibiotics. After growth overnight at 37 °C, colonies were picked into TB medium (Thermo Fisher Scientific) and incubated with shaking at 37 °C for 24 h. Cultures were collected using a QIAprep Spin Miniprep kit (Qiagen) according to the manufacturer’s instructions.

**Supplementary Text**

PsaCas12f mutations

To determine whether the model began to focus on specific locations of the protein, we further analyzed the pairwise mutational distance between the 12 mutations in each round and observed a shift to bimodality in the distribution of distances as the model entered the later rounds, with a bimodality coefficient of 0.72 in Round 4, suggesting that the model learned specific regions. We interpreted the mutations using a biophysical model to predict the free energy change of the protein based on the nominated mutations by our model. We found that the mutations nominated by the model in later rounds did not lead to protein stabilization (fig. S4H)(*13*). Instead, there was a large decrease in the coefficient of variation (COR) in the distribution of free energy changes, with only 471% COR in Round 4 relative to the 30,323% in Round 1 (fig. S4H). This selection runs contrary to traditional rational engineering approaches that try to maximize stability, and likely reflects the model's gain of understanding in the relationship between the fitness/stability and activity landscapes over iterative rounds, allowing the selection of non-intuitive residues. Protein activity is not necessarily correlated with stability, and thus the combination of the base LLM latent space model and the top layer "domain expert" model employed here represents an important step toward the efficient *in silico* evolution of higher activity protein variants.

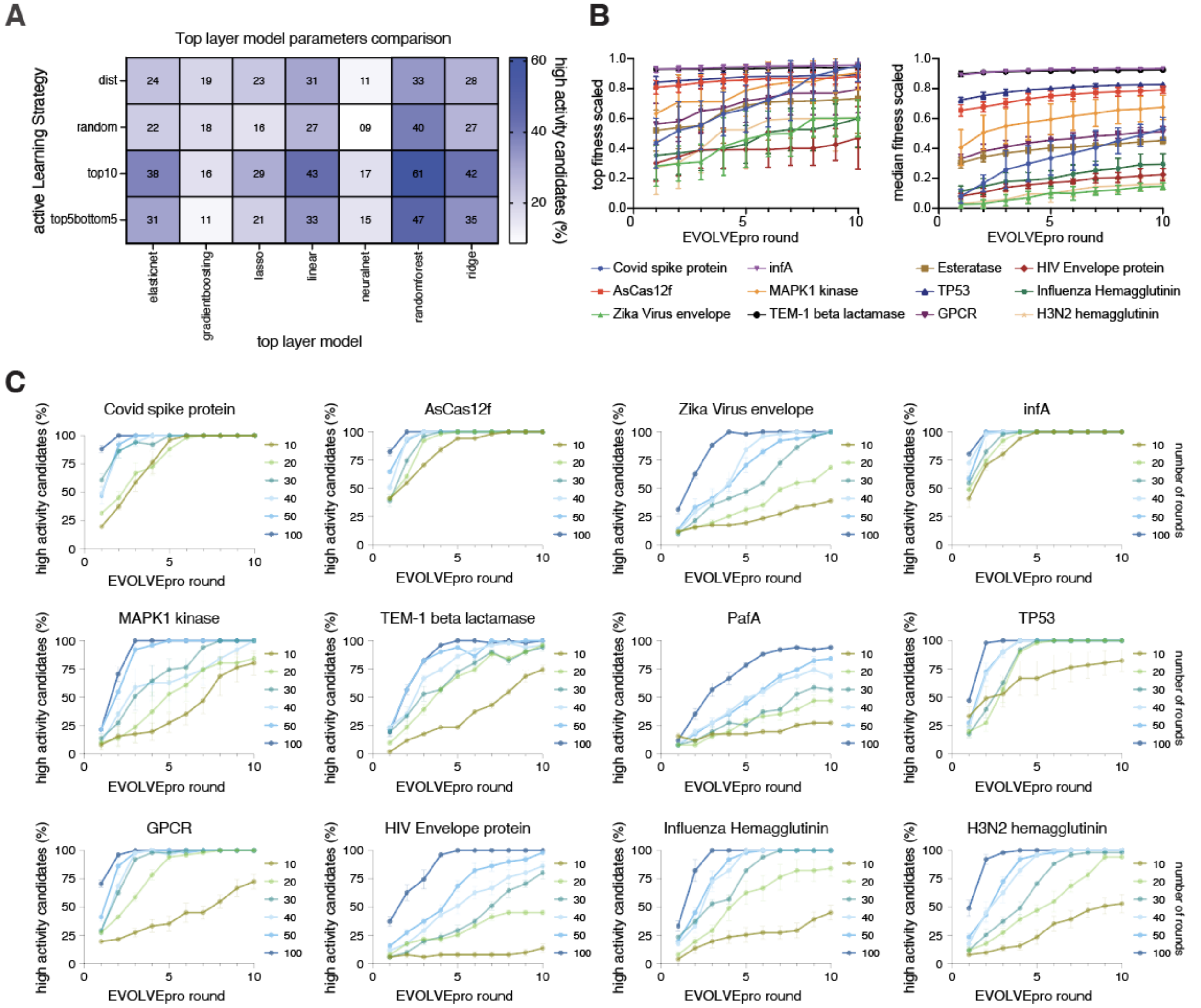


**Fig. S1. EVOLVEpro model optimization**

(**A**) Summary of parameter grid searches for EVOLVEpro with an ESM-2 15B foundational model, showing the random forest regressor combined with the top 10 active learning selection strategy returned the highest average binary top fitness success rate across 12 DMS datasets. (**B**) The optimized EVOLVEpro model with n=16 mutants per round shows both a higher median protein activity score and max protein activity score as the model progresses into later rounds of evolution, showing the utility of active learning. Error bars represent standard deviation with n=10 simulations. C) Comparison of the number of nominated mutants from n=10 to n=100 per round on the impact of EVOLVEpro evolution. Each line graph depicts the percent high fitness for one of the 12 DMS datasets and the error bar represents the standard error of the mean for 10 random simulations.

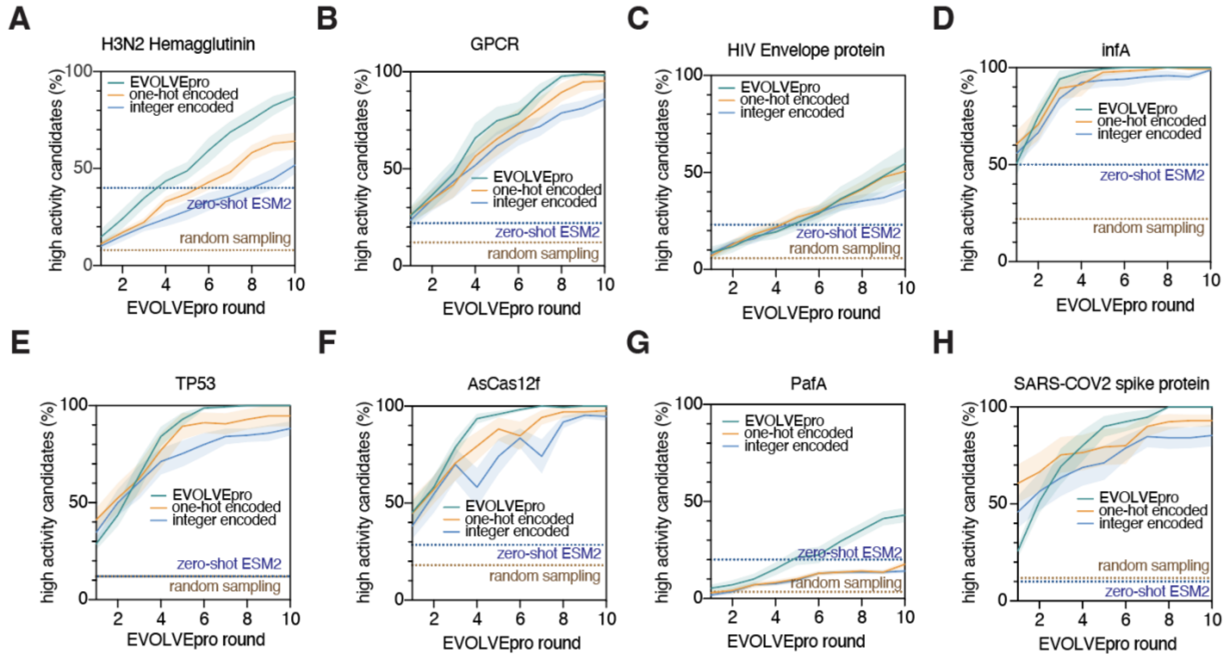


**Fig. S2. Additional EVOLVEpro model characterization**
(**A-H**) Performance over 10 rounds of EVOLVEpro with 16 mutants per round, compared to two different non-language model encoding schemes (One-hot encoding, integer encoding). Model performance is benchmarked on 8 additional datasets and compared to both the zero-shot ESM2 nomination success rate and background random sampling (*1*). Error bars represent the standard deviations for 10 random simulations.

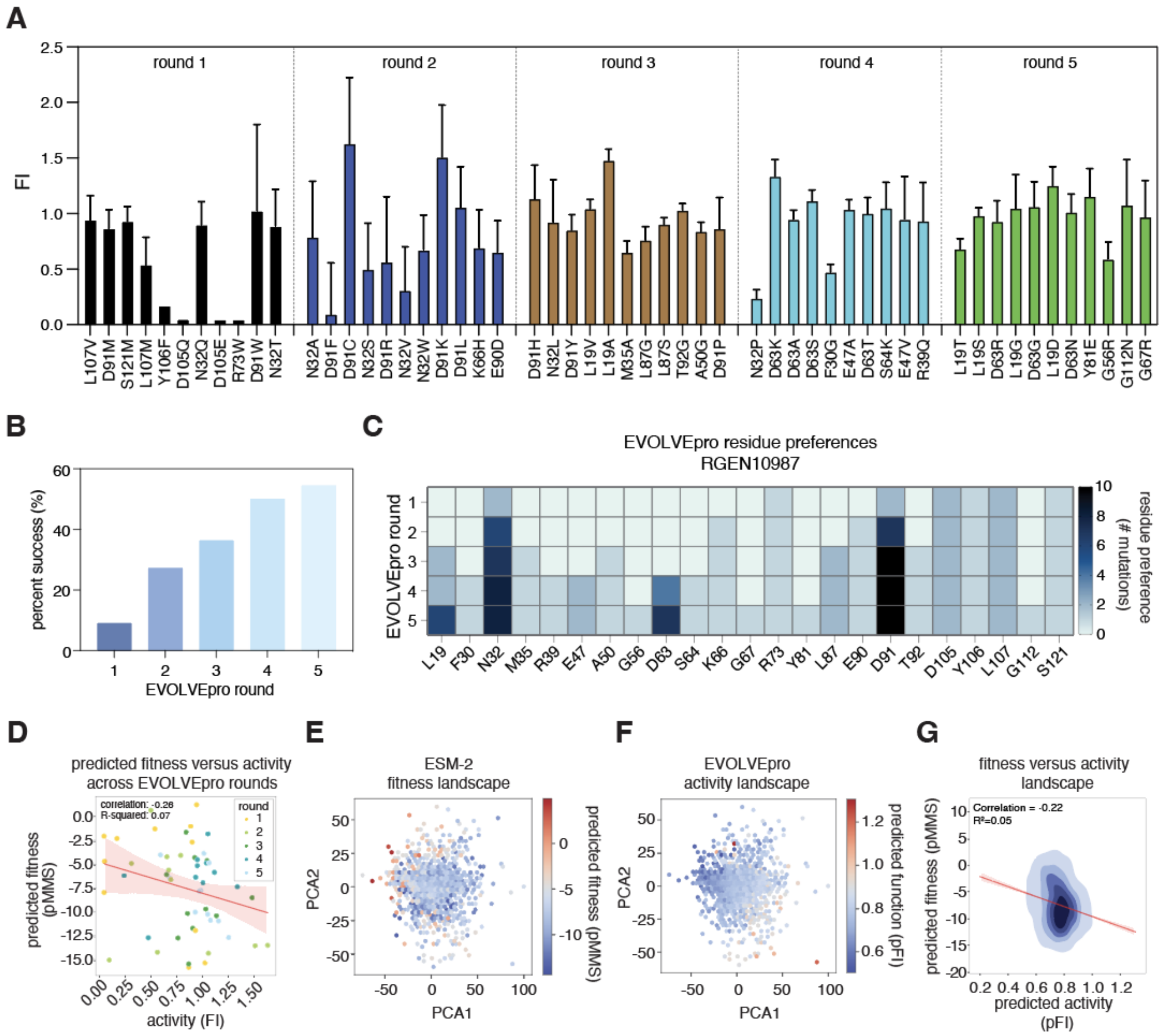


**Fig. S3. Additional characterization of evolution of REGN10987 with EVOLVEpro**

(**A**) Fold improvement of individual nominated REGN10987 mutants from EVOLVEpro across 5 rounds of evolution relative to WT. Error bars represent standard deviation of three biological replicates. (**B**) Percent success as defined by $IC_{50}$ lower than WT for the mutants in each round of evolution. (**C**) Heatmap showing the most common REGN10987 mutations explored by EVOLVEpro over rounds of evolution. Any position explored more than once is shown on a cumulative basis across rounds. (**D**) Scatter plot comparing the predicted ESM-2 protein fitness score versus experimentally measured REGN10987 binding affinity scaled fold improvement across evolution rounds. The correlation and linear regression line are shown in the plot. (**E-F**) Comparison of the REGN10987 antibody latent space with either predicted ESM-2 protein fitness (masked marginal score) or EVOLVEpro protein activity fold improvement. (**G**) A kernel density estimates of protein fitness as predicted by ESM-2 versus protein activity as predicted by EVOLVE-Pro. The correlation and linear regression line are shown in red, and the R square of the correlation is reported.

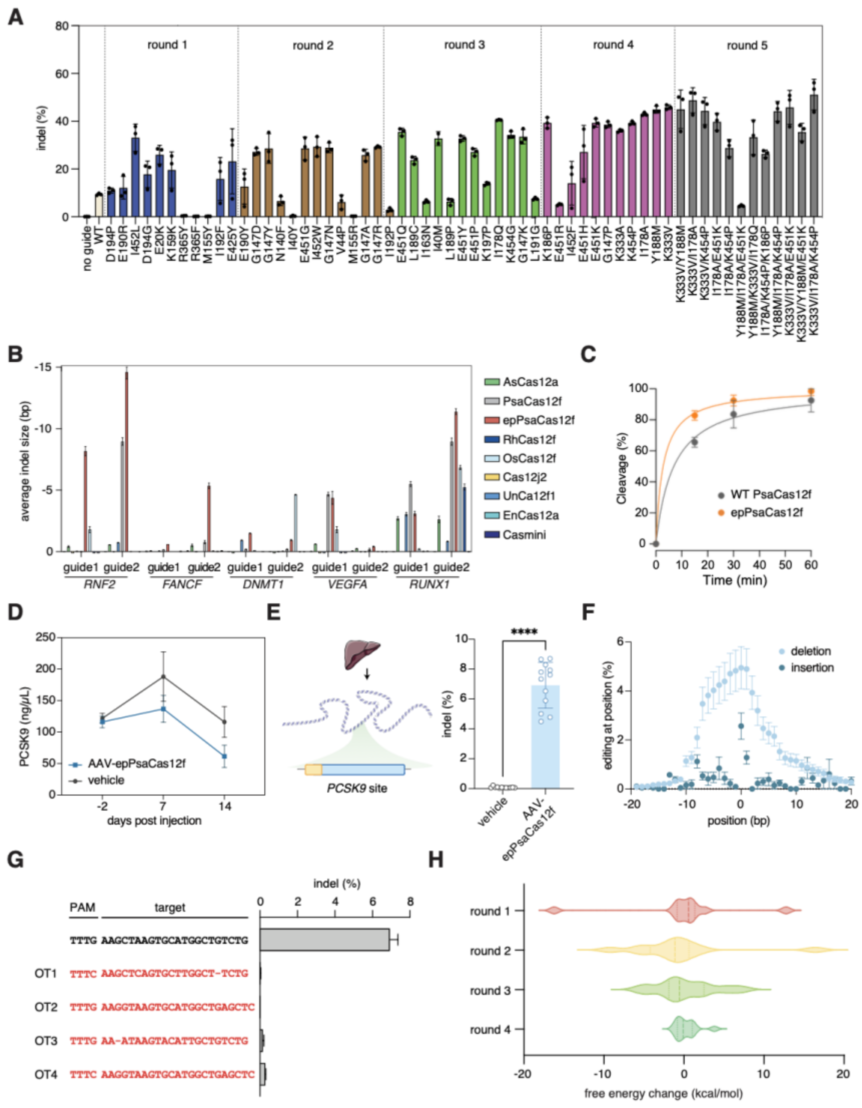


**Fig. S4. Additional characterization of evolution of PsaCas12f with EVOLVE-Pro**
(**A**) Indel editing rate of individual nominated mutants from EVOLVEpro across 5 rounds of evolution relative to WT. Error bars represent standard deviation of three biological replicates. (**B**) Average indel size generated by PsaCas12f, epPsaCas12f, and other Cas12f and Cas12a systems across 10 tested genomic regions. Error bars represent standard deviation of three biological replicates. (**C**) Kinetics of In vitro cleavage of a target DNA by either enPsaCas12 or WT PsaCas12f. Error bars represent standard deviation of three biological replicates. (**D**) Serum PCSK9 level determined by ELISA at -2, 7-, and 14-days post-injection of AAV-epPsaCas12f or a PBS control group. Error bars represent standard deviation of three biological replicates. (**E**) Next-generation sequencing determined editing in the livers of either AAV-epPsaCas12f or PBS-treated mice at the targeted PCSK9 site. Error bars represent standard deviation of 12 technical replicates. (**F**) Individual insertion and deletion frequency around the target spacer region in PCSK9 as quantified by CRISPREsso2. Error bars represent standard deviation of three biological replicates. (**G**) Specificity of epPsaCas12f. Indel formation was examined on the top four off-target (OT) sites predicted by Cas-OFFinder for epPsaCas12f with its PCSK9 targeting guide, showing minimal off-target editing at these OT sites(*14*). Error bars represent

standard deviation of three biological replicates. (**H**) A violin plot of free energy changes for each mutant in evolution compared to WT grouped by evolution round.

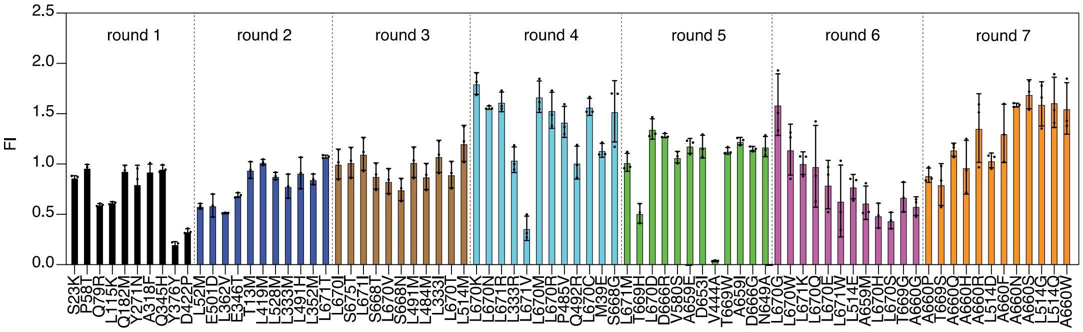


**Fig. S5. Additional characterization of the evolution of the prime editor PE2 with EVOLVE-Pro**

(**A**) Bar chart of each Individual PE2 mutant's fold improvement for murine genomic NOLC1 attB insertion frequency across evolution rounds. Error bars represent standard deviation of three biological replicates.

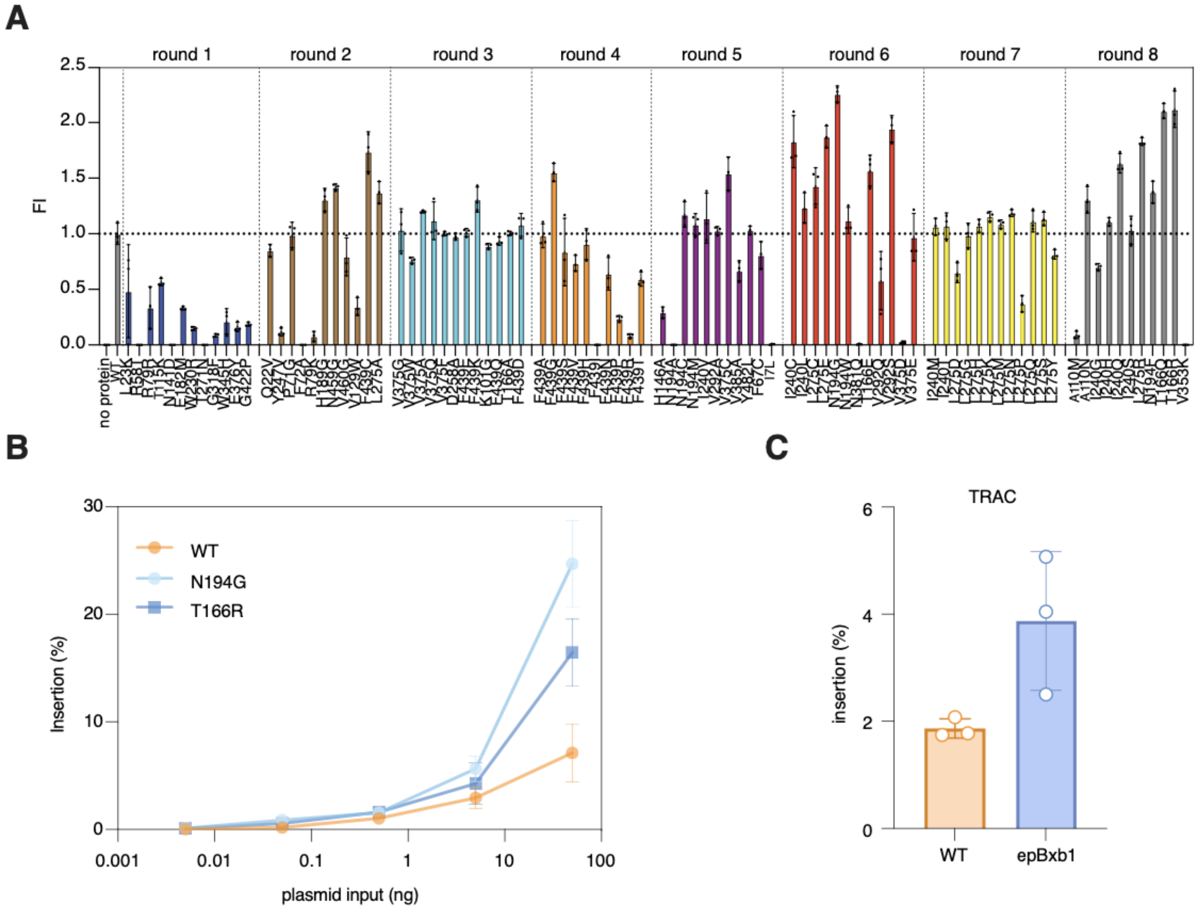


**Fig. S6. Additional characterization of the evolution of the Bxb1 integrase with EVOLVE-Pro**

(**A**) Individual mutant's fold improvement in plasmid recombination efficiency in HEK293FT cells across 8 rounds of evolution. (**B**) Titration of input Bxb1 amount for WT, N194G, and T166R (epBxb1) mutants in a genome integration assay in HEK293FT cells with attB sites preinstalled in the genome by lentivirus. Error bars represent standard deviation of three biological replicates. (**C**) Comparison of epBxb1 with wild-type Bxb1 on integration of cargo into TRAC genomic locus in HEK293FT cells. Error bars represent standard deviation of three biological replicates.

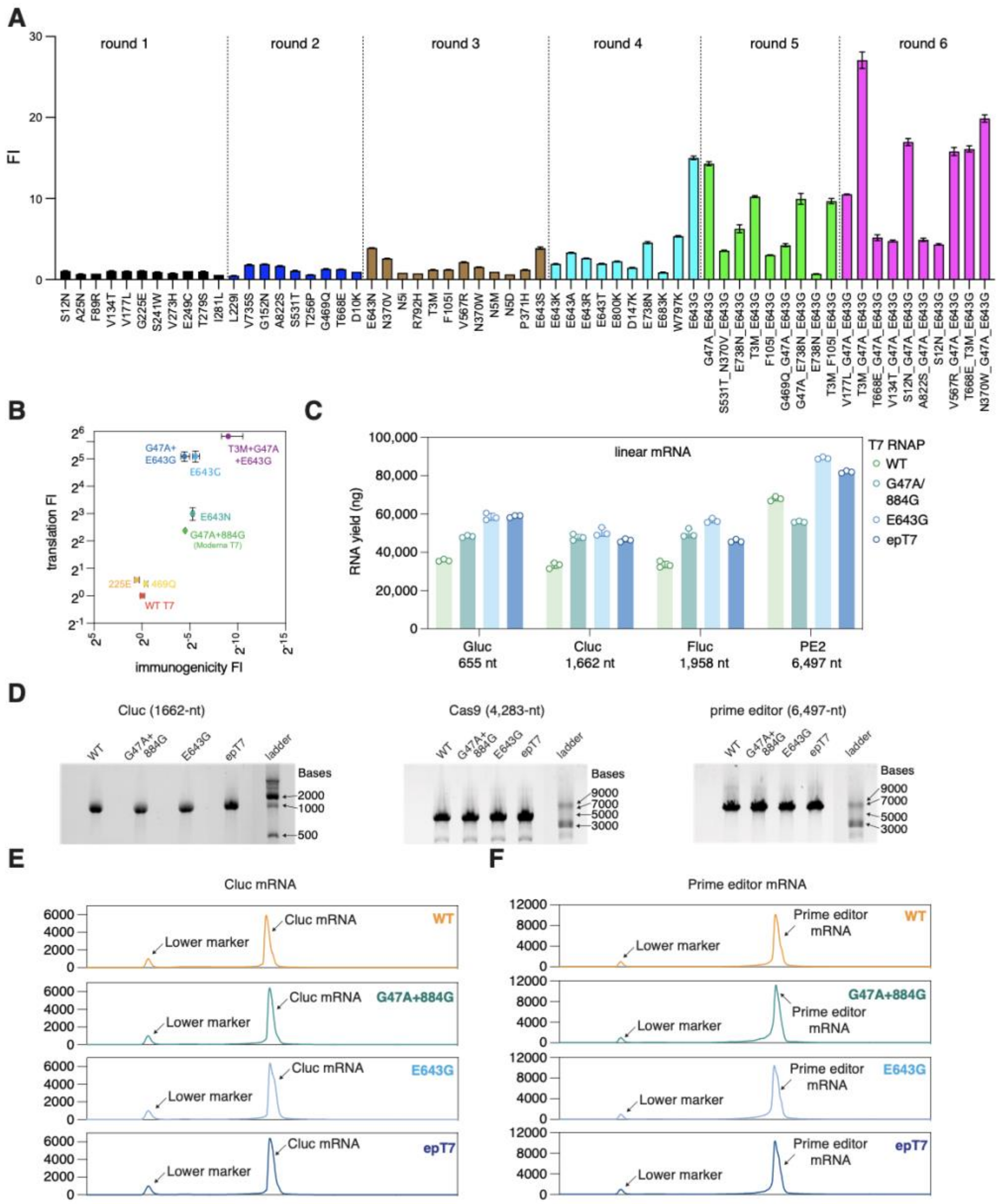


**Fig. S7. Characterization of evolution of T7 RNAP with EVOLVEpro**
(**A**) Individual mutant's fold improvement in transcription fidelity in producing Cypridina luciferase (Cluc) mRNA across 6 rounds of evolution. Error bars represent standard deviation of three biological replicates. (**B**) Comparison of each round's top-performing mutant with G47A/884G (previous SOTA) T7 RNAP in their production of Cluc mRNA. Two metrics are evaluated by assaying the protein translation and IFNB1 gene expression in BJ fibroblasts. Fold improvement is scaled to WT. (**C**) Comparison of the yield of mRNA across 4 different template sequences between WT, G47A/884G, E643G, and epT7. Error bars represent standard deviation of three technical replicates. (**D**) Gel electrophoresis analysis of 3 different mRNAs produced by different T7 RNAP variants: WT, G47A/884G, E643G, and epT7. (**E**) TapeStation analysis of Cluc mRNA produced by different T7 RNAP variants: WT, G47A/884G, E643G, and epT7. (**F**) TapeStation analysis of prime editor mRNA produced by different T7 RNAP variants: WT, G47A/884G, E643G, and epT7.

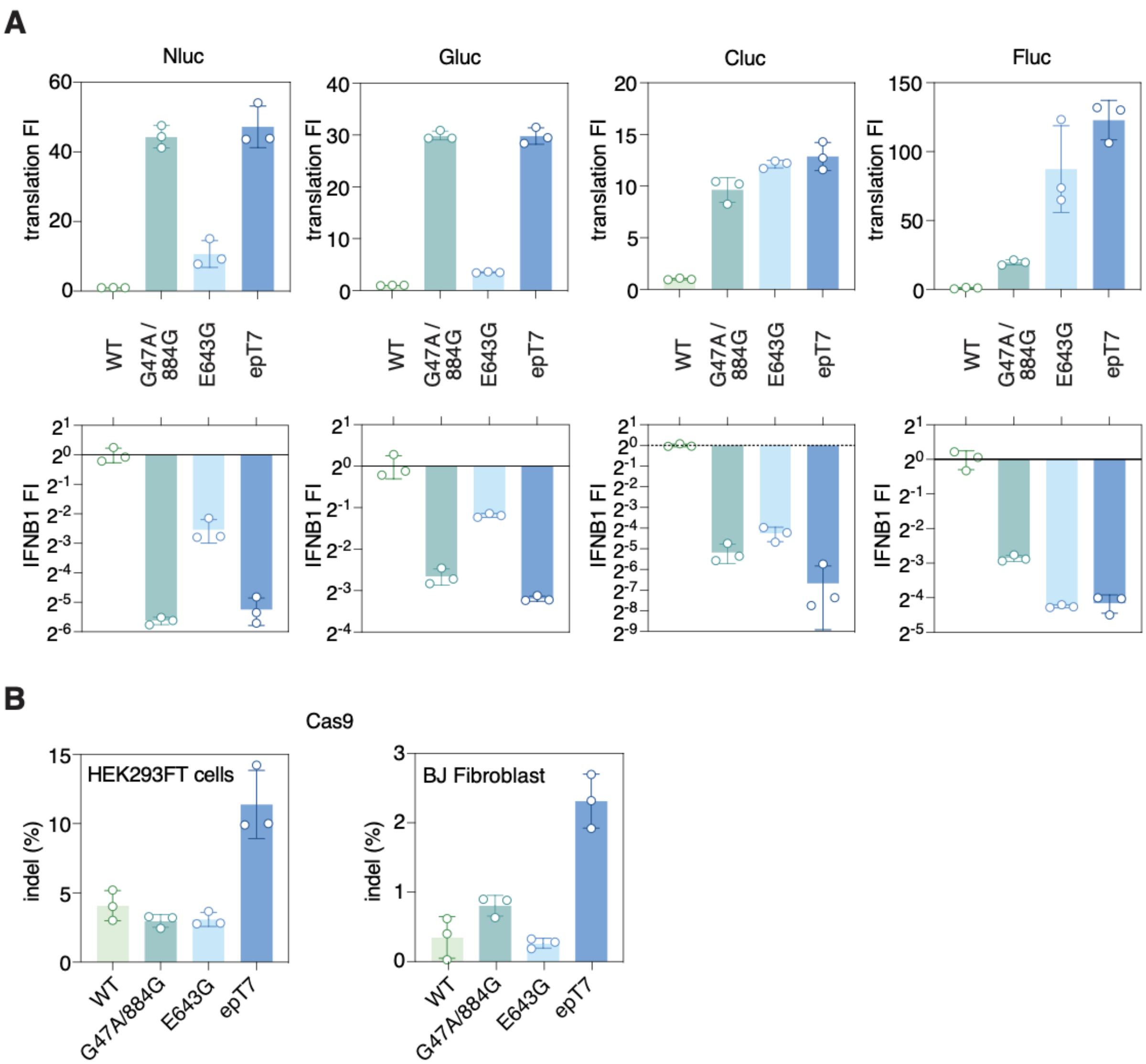


**Fig. S8. Additional characterization of the evolution of epT7 RNAP by EVOLVEpro**
(**A**) Translation and immunogenicity comparison of the previously engineered G47A/884G SOTA T7 RNAP with WT, E643G, and epT7 across 4 different template sequences in BJ Fibroblast cells. mRNAs were transfected into BJ fibroblast cells for either protein translation readout or targeted IFNB1 gene expression analysis using qPCR 24 hours after transfection. Error bars represent standard deviation of three biological replicates. (**B**) Comparison of editing activity of Cas9 mRNA produced by WT, G47A/884G, E643G, and epT7 on endogenous ENO1 genomic loci in HEK293FT cells and BJ Fibroblast cells. Indel rates are quantified 48 hours after transfection of Cas9 mRNA and synthetic guide RNA targeting ENO1. Error bars represent standard deviation of three biological replicates.

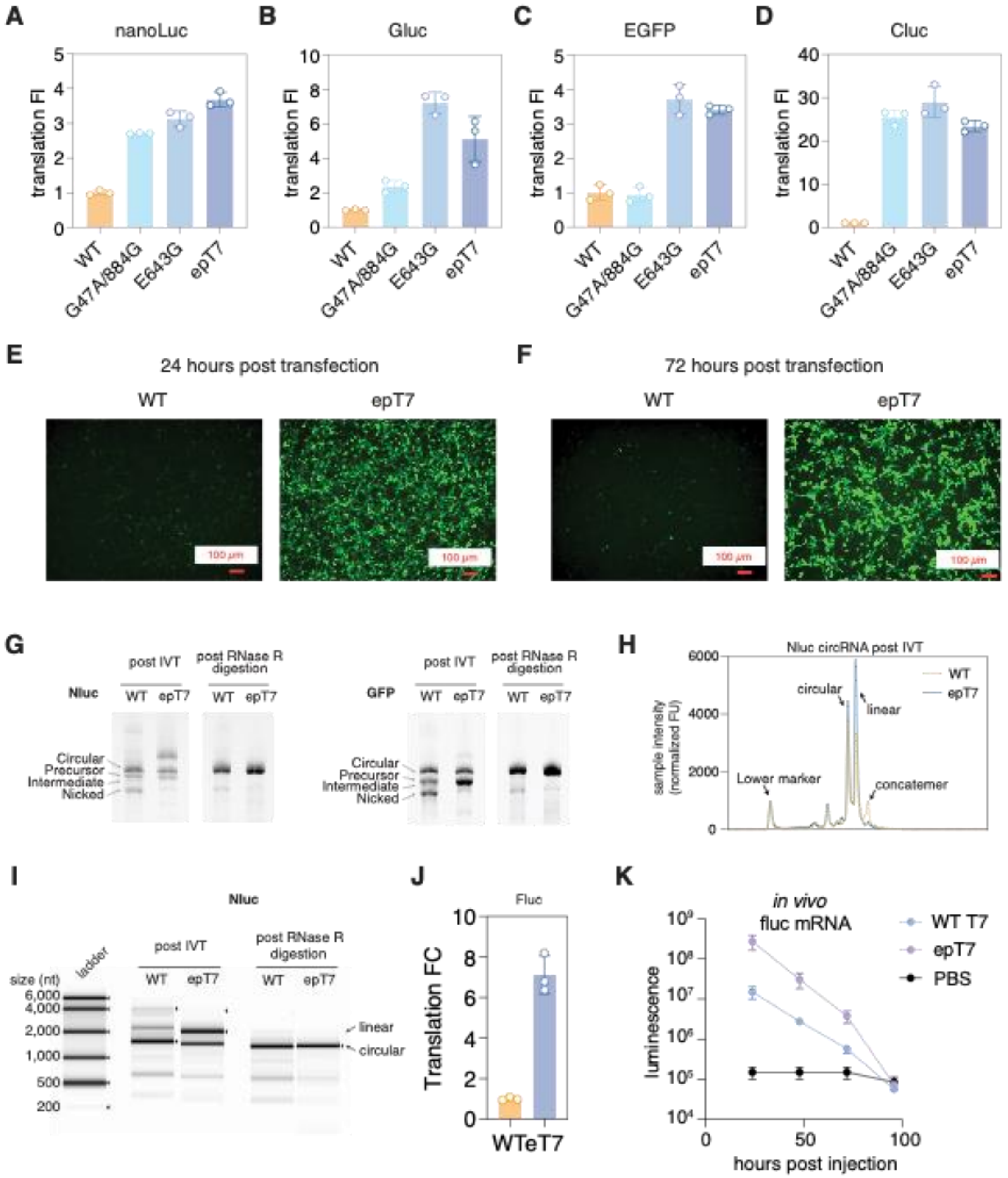


**Fig. S9. Circular RNA characterization and *in vivo* bioluminescence of RNAs produced by epT7 RNAP**

(**A-D**) Validation of epT7 produced circRNA sequences compared to G47A/884G, $T7^{E643G}$, and wild-type T7 on A) Nanoluc, B) Gluc, C) EGFP, D) Cluc encoded circRNAs. Translation of each protein is measured in HEK293FT cells 48 hours after transfection. (**E-F**) Representative fluorescence images for circular GFP RNA produced by wild-type T7 RNAP or epT7 at E) 24 hours and F) 72 hours post-transfection in HEK293FT cells. Scale bar, 100 µm. (**G**) Migration of Nanoluc and GFP encoded circRNAs produced by either wild-type T7 or epT7 on E-Gel EX systems. (**H-I**) TapeStation analysis of pre and post-RNaseR cleaned-up nanoluc circular RNA produced by either wild-type T7 or epT7. (**J**) Validation of epT7 produced circRNA on firefly luciferase sequences compared to wild-type T7. Translation of each protein is measured in HEK293FT cells 48 hours after transfection. Error bars represent standard deviation with n=3 biological replicates. (**K**) Time course kinetics of firefly luciferase luminescence (log scale) at 4 different time points after injection of LNP encapsulated mRNA made by either wild-type T7 or epT7 compared to background PBS control. Error bars represent standard deviation with n=3 biological replicates.

### Table S1. Summary of parameters grid search

| Fitness measurement data type | First-Round strategy | Active learning strategy | PLM embedding types | Top layer model | Round Number | Mutant Nomination numbers |
|---|---|---|---|---|---|---|
| Raw input | Random | Random | Raw | Ridge regression | 5 | 10 |
| Min-Max scaled input | Max distance K-medoids clusters | Top N | PCA (Top 10 PCs) | Lasso regression | 10 | 20 |
| | | Top N/2 + Bottom N/2 | Residue Averaged | Elastic Net | | 30 |
| | | Max Euclidean distance | | Multilayer perceptron | | 40 |
| | | | | Linear Regression | | 50 |
| | | | | Neural Network | | 100 |
| | | | | Random Forest Regressor | | 200 |
| | | | | XGboost | | 500 |

### Table S2. Description for 12 DMS datasets

| **Protein** | **Reference** | **Organism** | **Fitness setting** | **Cutoff** | **Population successes** | **Population size** | **Efficient evolution rate (hie et al 2023)** | **Background (%)** |
|---|---|---|---|---|---|---|---|---|
| ADRB2 | Jones et al., 2019 | Human | Signal transduction + pathway reporter | > 2.8 | 914 | 7800 | 22 | 12 |

| | | | | | | | | |
|---|---|---|---|---|---|---|---|---|
| β-lactamase | Stiffler et al, 2015 | Bacteria | Antibiotic resistance (ampicillin, 2500 ug/mL) | > 0.01 | 393 | 4978 | 40 | 7.9 |
| Env | Haddox et al., 2016 | Virus | Viral replication fitness | > 0.1 | 748 | 12863 | 23 | 5.8 |
| HA H1 | Doud and Bloom, 2016 | Virus | Viral replication fitness | > 0.1 | 645 | 10716 | 16 | 6 |
| HA H3 | Lee et al., 2018 | Virus | Viral replication fitness | > 0.1 | 714 | 10754 | 31 | 6.6 |
| infA | Kelsic et al., 2016 | Bacteria | Competitive growth | > 0.98 | 305 | 1368 | 50 | 22 |
| MAPK1 | Brenan et al., 2016 | Human | Competitive growth (SCH772984) | > 2.5 | 77 | 6810 | 7.7 | 1.1 |
| P53 | Giacomelli et al, 2018 | Human | Competitive growth (etoposide) | > 1 | 905 | 7448 | 12 | 12 |
| PafA | Markin et al., 2021 | Bacteria | Kcat/KM | P < 0.01, faster than WT | 35 | 1040 | 20 | 3.4 |
| AsCas12f | Hino et al., 2023 | Bacteria | Genomic DNA Clevage | > 1 | 1436 | 7941 | 28 | 18 |
| Zika Envelope protein | Sourisseau et al., 2019 | Virus | Viral replication fitness | > 1 | 351 | 9576 | 12 | 3.7 |
| COV2 Spike Protein | Greaney et al., 2021 | Virus | Yeast display binding | > 0.05 | 232 | 1959 | 0 | 12 |

**Data S1. EVOLVEpro parameter grid search results**
**Data S2. Raw data for 12 DMS datasets**
**Data S3. EVOLVEpro protein language model comparison**
**Data S4. EVOLVEpro zero-shots training comparison**
**Data S5. Guide sequences used in this study**
**Data S6. Next-generation sequencing primer sequences used in this study**
